# Nitrogenase regulation in *Vibrio natriegens* differs from other γ-proteobacteria

**DOI:** 10.64898/2026.05.30.728972

**Authors:** Nicholas W. Haas, Emma E. Wiesler, Ankur B. Dalia, Xindan Wang, James B. McKinlay

## Abstract

Regulation of nitrogenase, which converts nitrogen gas (N_2_) into ammonium (NH_4_^+^), typically involves a conserved set of regulatory proteins across diverse N_2_-fixing (diazotrophic) bacteria. However, the interactions and relative influence of these regulators can vary between species. Thus, one cannot make assumptions about nitrogenase regulation when working with uncharacterized diazotrophs like *Vibrio natriegens*, a γ-proteobacterium of growing interest for synthetic biology. Little is known about *V. natriegens* nitrogenase regulation, which could be used to exploit inexpensive N_2_ for various applications, including NH_4_^+^ production. Here, we characterized the roles of several annotated *V. natriegens* nitrogenase regulatory proteins in response to NH_4_^+^ versus N_2_. Using functional genomics, targeted mutations, and reporter assays, we identified a typical regulatory hierarchy where the two-component system NtrBC governs a nitrogen-scavenging regulon that includes NifA, the transcriptional activator of nitrogenase genes. Unlike other diazotrophic γ-proteobacteria, NifA was sufficient to activate nitrogenase gene expression, as a mutant lacking NtrBC grew normally with N_2_ after a lag phase. Thus, NtrBC was dispensable, but still important for timely nitrogenase expression. Furthermore, NtrBC was negatively regulated by the nitrogen-responsive P_II_ proteins GlnB and GlnK; disruption of both P_II_ proteins led to NtrBC-dependent nitrogenase overactivity, marked by NH_4_^+^ excretion. The redundant repression of NtrBC by GlnB and GlnK more closely resembles that of non-diazotrophic *E. coli* than other diazotrophic γ-proteobacteria. Together, our findings provide a framework for *V. natriegens* nitrogenase regulation that can be leveraged for applications like NH_4_^+^ production.

**HIGHLIGHTS:**

- A genetic examination of *Vibrio natriegens* nitrogenase regulation is performed
- NtrBC is important for early nitrogenase gene expression but is not essential
- NifA autoactivation is sufficient for nitrogenase expression
- P_II_ proteins GlnB and GlnK are redundant negative regulators of nitrogenase
- Genetic targets are identified that result in excretion of NH_4_^+^

**GRAPHICAL ABSTRACT:** 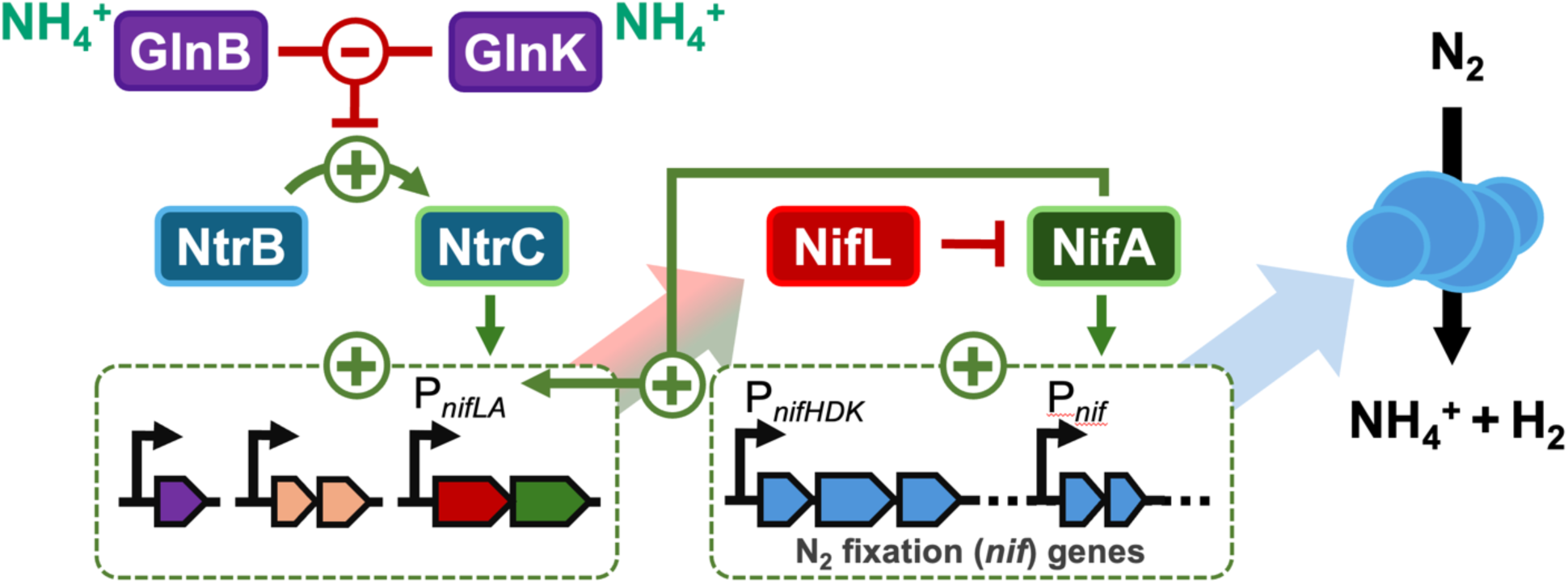

## INTRODUCTION

Nitrogenase is a critical enzyme that sustains life by converting abundant nitrogen gas (N_2_) into ammonium (NH_4_^+^) in a process called N_2_-fixation or diazotrophy. Only certain bacteria and archaea are diazotrophs, and their combined activity represents a major input of nitrogen into ecosystems [1]. Currently, the largest terrestrial nitrogen input comes from synthetic agricultural fertilizers produced by the industrial Haber-Bosch process, which is energy-intensive, polluting, and inaccessibly expensive for some developing regions [1–3]. Despite these undesirable aspects, Haber-Bosch fertilizers are important to sustain half the world’s population [3]. Thus, there are incentives to produce clean and affordable alternative fertilizers by harnessing diazotrophs [4, 5]. To do so requires an understanding of how diazotrophs regulate nitrogenase in response to environmental signals like NH_4_^+^. Nitrogenase regulation is well-defined for several diazotrophs, but other diazotrophs of emerging biotechnological interest, like the marine γ-proteobacterium *Vibrio natriegens* ATCC 14048, have not been characterized.

*V. natriegens* has become popular in the last decade as a chassis for synthetic biology due to its rapid growth and metabolism [6–8] and its genetic tractability [9–12]. As a diazotroph, *V. natriegens* can also use N_2_ as the sole nitrogen source under anoxic conditions [13]. Accessing inexpensive N_2_ could provide cost-savings for industrial fermentations [14], including for NH_4_^+^ production. However, since first being described in 1988, *V. natriegens* N_2_ fixation has received little attention beyond measuring responses in nitrogenase structural *nifHDK* gene expression to fixed nitrogen (e.g., NH_4_^+^, peptone, etc) and oxygen (O_2_) [15] and verification of functional activity from a predicted nitrogenase gene cluster [16]. Thus, it is largely unknown how *V. natriegens* nitrogenase is regulated. Moreover, little is known about nitrogenase regulation for any *Vibrio* species beyond a recent investigation with *V. diazotrophicus* [17], and for marine heterotrophic diazotrophs in general, despite their abundance and importance in marine ecosystems [18, 19].

Regulation of nitrogenase typically involves multiple interacting regulatory modules, ensuring tight control over its energetically expensive production and operation; at least 20 genes are required to assemble nitrogenase, which then requires 16 ATP and 8 electrons per 2 NH_4_^+^ produced (Eq 1) [20]. In one of the best characterized γ-proteobacterial diazotrophs, *Klebsiella pneumoniae*, the two-component system NtrBC forms the first module, governing a diverse regulon in response to nitrogen starvation that includes nitrogenase and other N_2_-fixation genes (*nif* genes). When fixed nitrogen is available, the sensor kinase NtrB dephosphorylates the NtrC response regulator, preventing expression of nitrogen scavenging genes (Fig 1A, left) [20]. As fixed nitrogen becomes limiting, NtrB phosphorylates NtrC, which then serves as a transcriptional activator of nitrogen scavenging genes [20] (Fig 1A, right).

**Fig 1.**
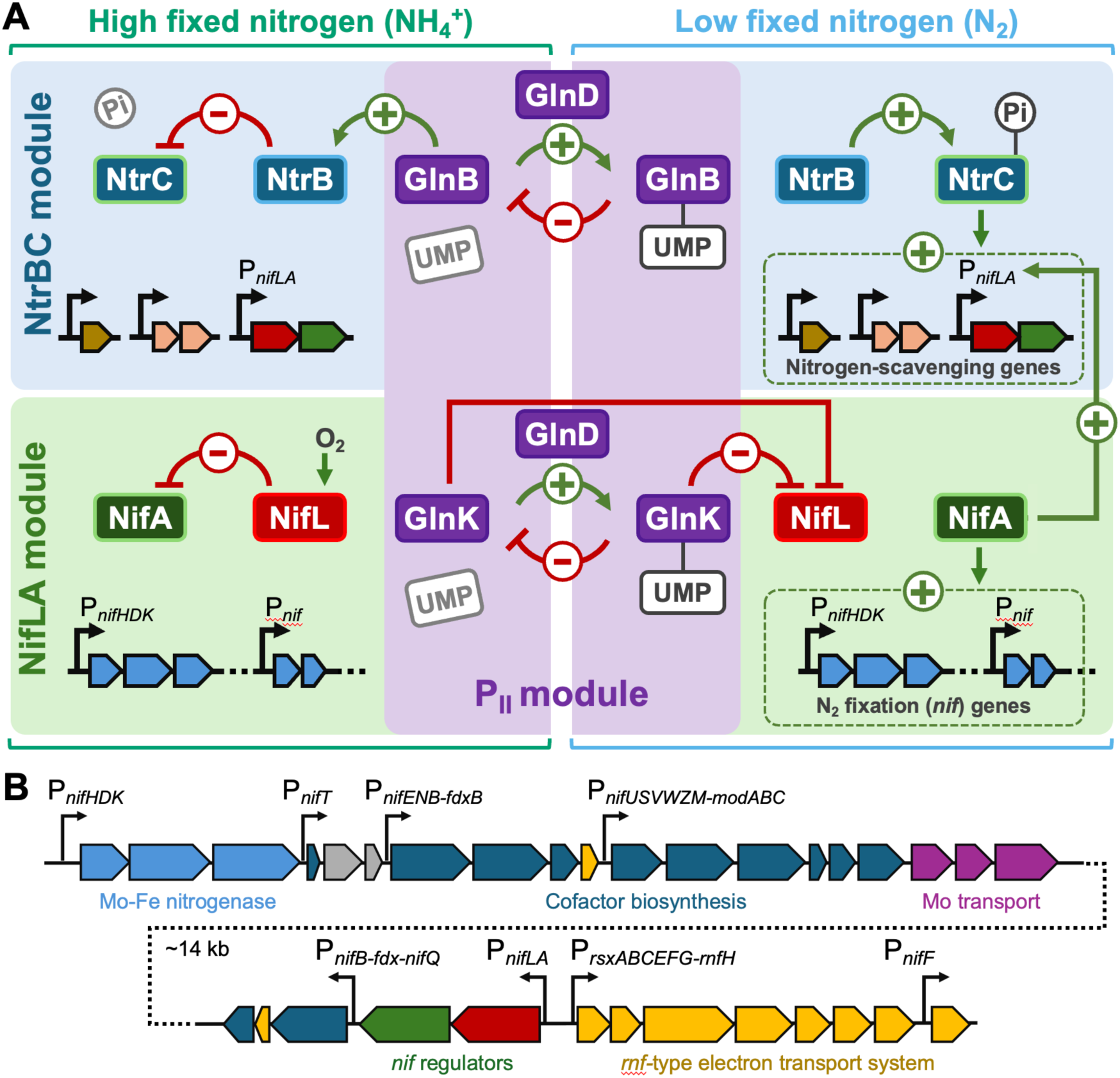
Nitrogenase regulation in *K. pneumoniae* and the *V. natriegens* nitrogenase gene clusters. **(A)** Nitrogenase regulation in the γ-proteobacterium, *K*. *pneumoniae*, as a comparative reference for characterizing *V. natriegens* nitrogenase regulation, which has a similar inventory. There are three regulatory modules: (i) the two-component system NtrBC, which activates a regulon of nitrogen scavenging genes in response to low fixed nitrogen availability, (ii) NifLA, where the anti-activator NifL determines the availability of NifA to directly activate transcription of nitrogenase genes (*nif* genes), and (iii) P_II_-proteins GlnB and GlnK, which sense fixed nitrogen availability and respond by influencing the NtrBC activity (via GlnB) or NifL-NifA interactions (via GlnK). P_II_-activity is mediated by post-translation modification with uridine monophosphate (UMP) via GlnD (Fig 1A) [20]. NifL-NifA interactions are also influenced by O_2_ sensing via NifL. **(B)** Organization of *nif* genes on *V. natriegens* ATCC 14048 chromosome 2.

Among these genes is *nifA*, which encodes the master transcriptional activator of *nif* genes. In *K. pneumoniae*, *nifA* is co-transcribed with *nifL*, encoding an anti-activator that represses NifA in a redox-responsive manner [20–23]. NifLA thus forms the 2nd regulatory module (Fig 1A). Finally, a third module of P_II_ proteins GlnB and GlnK modulates the activity for the other tiers in response to fixed nitrogen availability (Fig 1A). In *K. pneumoniae*, GlnB is constitutively expressed and modulates NtrB activity [24]. GlnK, expressed under the control of NtrBC, relieves NifL inhibition of NifA when fixed nitrogen is scarce [25–27]. P_II_ protein activity can also be modulated by the addition or removal of UMP groups by GlnD [20]. However, in *K. pneumoniae*, GlnK modification with UMP does not affect its activity on NifL [25, 28, 29] (Fig 1A). Like *K. pneumoniae*, *V. natriegens* ATCC 14048 does not appear to encode a fourth module of post-translational regulation involving DraTG nor does it encode either alternative nitrogenase [20] (Fig 1B, Table S1).

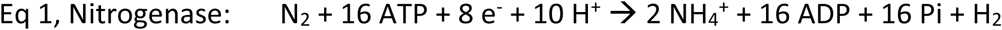

Whereas the above regulatory proteins are commonly found across diverse diazotrophs, they can interact in different ways and exert different levels of control. For example, three different purple phototrophic bacteria with similar inventories of nitrogenase regulatory proteins required mutations in different regulatory proteins to achieve constitutive nitrogenase activity [30–33]. Even for *V. diazotrophicus*, where *nif* expression was shown to be NtrC-dependent [34], there is little shared synteny in the *nif* gene clusters with *V. natriegens*, suggesting different evolutionary histories and possibly distinct mechanisms of *nif* regulation [16, 35]. Thus, the influence that each regulatory protein has over nitrogenase cannot be reliably inferred from a genome sequence; it must be interrogated directly.

Here, we provide the first genetic characterization of the *V. natriegens* nitrogenase regulatory hierarchy. We find that unlike other γ-proteobacterial diazotrophs, NtrBC is not required for N_2_ fixation. NtrBC facilitates early *nif* gene expression in response to low nitrogen, but wild-type diazotrophic growth rates can still be achieved without NtrBC. Instead, NifA autoactivation is sufficient for a high diazotrophic growth rate. Also differing from other diazotrophic γ-proteobacteria, *V. natriegens* P_II_ proteins play redundant roles in preventing unchecked nitrogenase activity via NtrBC. Mutants lacking both P_II_ proteins exhibited high nitrogenase activity, manifesting in NH_4_^+^ excretion. Our work provides a foundational framework of *V. natriegens* nitrogenase regulation and exposes engineering targets for NH_4_^+^ production.

## RESULTS

### Activation of NifA is required for *V. natriegens* nitrogenase activity

To characterize *V. natriegens* nitrogenase regulation, we first verified that *V. natriegens* fixes N_2_ when grown fermentatively in an anoxic minimal medium with glucose. Indeed, the diazotrophic growth rate (nitrogen source: N_2_) was 0.33 ± 0.05 h^-1^, 42% of the growth rate with NH_4_^+^ (0.79 ± 0.03 h^-1^; mean ± SD; Fig 2A, B; note different x-axis scales). We then addressed the essentiality of NifA, the canonical master transcriptional regulator of *nif* genes, for diazotrophic growth [20]. When we deleted *nifA*, the resulting Δ*nifA* mutant grew with NH_4_^+^ but not with N_2_ (Fig 2A, B). Thus, NifA is required for *V. natriegens* diazotrophic growth.

**Fig 2.**
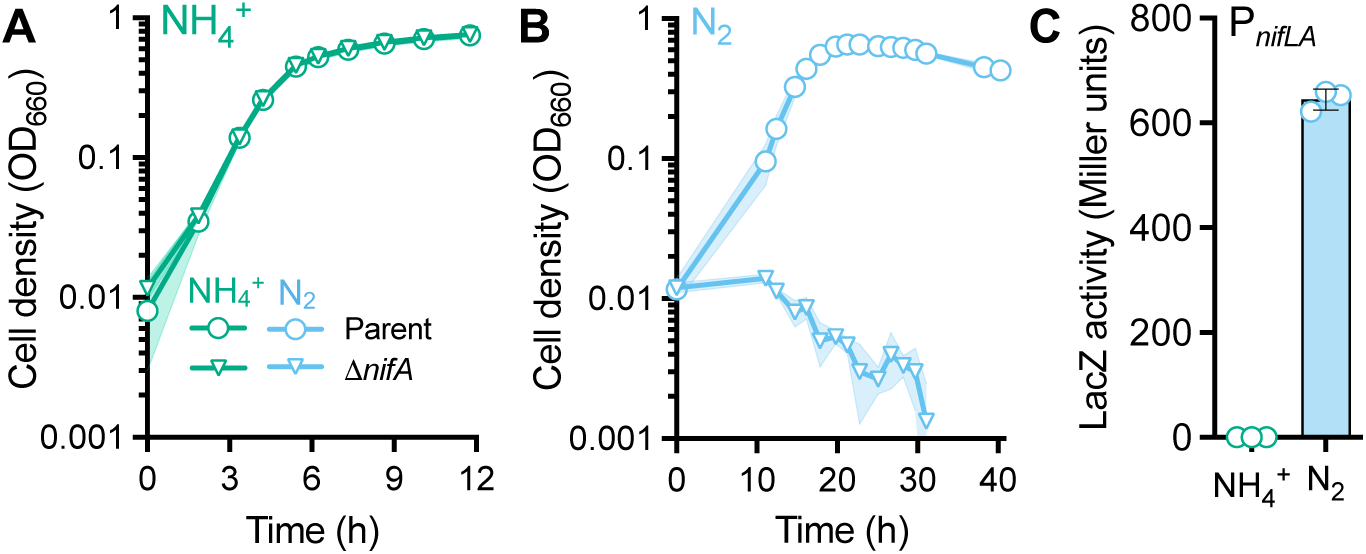
NifA is essential for growth on N_2_. **(A, B)** Growth curves for *V. natriegens* strains grown in anoxic minimal media with glucose and either NH_4_^+^ **(A)** or N_2_ **(B)** as the sole nitrogen source. Parent, NWH003 (Δ*dns*::Km^R^); Δ*nifA*, NWH036 (Δ*dns*::Km^R^). Points, mean; shading, SD; n=3. **(C)** A P*_nifLA_*-*lacZ* transcriptional reporter was used to compare *nifLA* expression in the parent (NWH010; Δ*dns*::Sp^R^*-*P*_nifLA_*-*lacZ*) when grown with NH_4_^+^ or N_2_. Points, biological replicates; bars, mean; error bars, SD; n=3.

*nifA* is often co-transcribed with *nifL*. NifL conventionally binds NifA and represses transcription in a redox-dependent manner [23]. In some γ-proteobacteria like *Azotobacter vinelandii*, *nifLA* is constitutively expressed [36, 37] whereas in *K. pneumoniae* and *Pseudomonas stutzeri*, *nifLA* is transcribed only when fixed nitrogen is scarce [38, 39]. To determine if *V. natriegens nifLA* expression was constitutive or responsive to NH_4_^+^ availability, we measured *nifLA* promoter (*P_nifLA_*) activity using a LacZ transcriptional reporter (P*_nifLA_*-*lacZ*) in WT *V. natriegens* grown with NH_4_^+^ versus N_2_. We only observed LacZ activity under N_2_-fixing conditions (Fig 2C). Thus, *V. natriegens nifLA* expression is responsive to NH_4_^+^ availability.

NifL typically modulates nitrogenase activity by inhibiting NifA through protein-protein interactions in response to fixed nitrogen or O_2_ [40]. To determine the extent to which NifL controls nitrogenase activity in *V. natriegens*, we first compared growth between the parent and a Δ*nifL* mutant during anaerobic growth with NH_4_^+^ versus N_2_. Growth trends were superimposable with NH_4_^+^, whereas with N_2_ the Δ*nifL* mutant exhibited a similar growth rate but a lower final cell density compared to the parent (Fig 3A). Based on these growth trends, we predicted that (i) *nif* genes would not be expressed with NH_4_^+^ in the Δ*nifL* mutant and (ii) that there might be excessive nitrogenase expression with N_2_, creating an energetic burden. Indeed, P*_nifLA_-lacZ* reporter activity was not observed in either the parent or the Δ*nifL* mutant when grown with NH_4_^+^. With N_2_, P*_nifLA_-lacZ* activity was 2-fold higher in the Δ*nifL* mutant compared to the parent (Fig 3B). This increase could be due to NifA, no longer inhibited by NifL, increasing *nifA* expression (autoactivation), which can occur in other diazotrophs like *K. pneumoniae*, according to reporter and biochemical assays [41–43]; we directly test this possibility later in the study. Similarly, when we measured expression of the nitrogenase structural genes using a P*_nifHDK_*-*lacZ* reporter under diazotrophic conditions, LacZ activity was 1.7-fold higher in the Δ*nifL* mutant compared to the parent (Fig 3C). This increased *nif* gene expression translated to increased nitrogenase activity, measured as H_2_ production, an obligate coproduct of nitrogenase activity (Eq 1); we previously verified that *V. natriegens* does not have hydrogenases to make H_2_ and it does not produce H_2_ when supplied with NH_4_^+^, conditions where nitrogenase is repressed [44]. H_2_ production from the Δ*nifL* mutant was 1.4-fold higher than the parent (Fig 3D). These data indicate that NifL alone does not control NifA expression or activity in response to NH_4_^+^, but it might have a dampening effect on *nif* transcription under N_2_-fixing conditions, where we infer that some NifA is always bound and inhibited by NifL.

**Fig 3.**
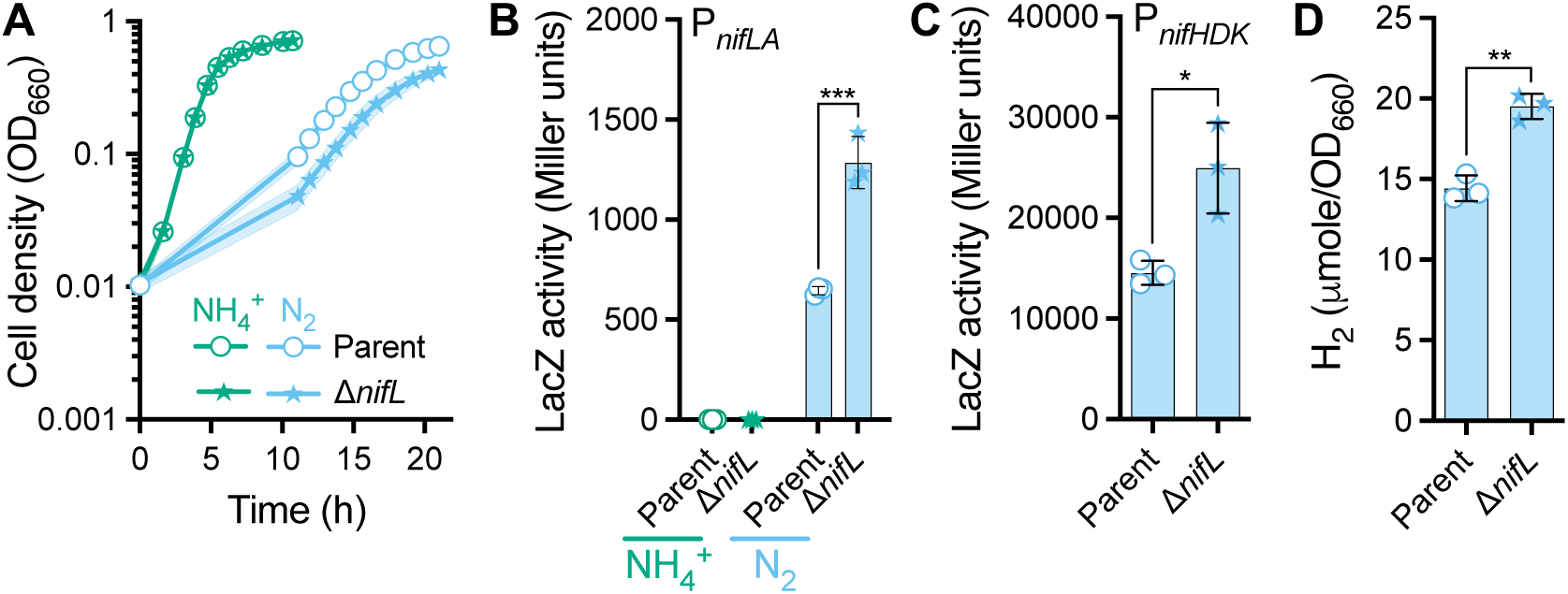
NifL is a negative regulator of nitrogenase expression and activity. **(A)** Growth curves for *V. natriegens* strains grown in anoxic minimal media with glucose and either NH_4_^+^ or N_2_ as the sole nitrogen source. Parent, NWH003 (Δ*dns*::Km^R^); Δ*nifL*, NWH037 (Δ*dns*::Km^R^). Points, mean; shading, SD; n=3. **(B, C)** A *lacZ* transcriptional reporter was used to compare expression from P*_nifLA_* **(B)** or P*_nifHDK_* **(C)** under N_2_-fixing conditions. **(D)** Nitrogenase activity was measured in stationary phase (13 h) as accumulated H_2_ (Eq 1). **(B-D)** Points, biological replicates; bars, mean; error bars, SD; n=3. Statistical differences were determined using unpaired, two-tailed t tests; *, *P* < 0.05; **, *P* < 0.01; ***, *P* < 0.001.

Although herein we primarily focus on nitrogenase regulation in response to NH_4_^+^ versus N_2_, we also verified whether NifL prevents *nif* gene expression in response to O_2_. To induce nitrogenase expression during aerobic growth, we used glutamate as the sole nitrogen source. Glutamate can ‘trick’ some diazotrophs into expressing nitrogenase even though glutamate is available as a nitrogen source [45]. *V. natriegens* grew aerobically with glutamate, albeit with a growth rate that was 15% of that with NH_4_^+^ (Fig S1A). To test whether glutamate induced *nif* gene expression, we first measured P*_nifLA_*-*lacZ* reporter activity. Indeed, glutamate led to *nifLA* expression in the parent (Fig S1B). The Δ*nifL* mutant had P*_nifLA_*-*lacZ* activity that was 2.7-fold higher than the parent (Fig S1B), again suggesting possible NifA autoactivation. We then determined if NifL ultimately prevents nitrogenase expression during aerobic growth by measuring P*_nifHDK_*-*lacZ* reporter activity. Indeed, despite the expression from P*_nifLA_*, no P*_nifHDK_*-*lacZ* activity was observed in the parent. The Δ*nifL* mutant P*_nifHDK_*-*lacZ* activity was about as high as Δ*nifL* mutant levels under N_2_-fixing conditions (Fig. S1C vs Fig 3C). Thus, NifL prevents nitrogenase expression in response to O_2_.

### NtrBC activates *nifLA* expression but is dispensable for N_2_ fixation

The lack of P*_nifLA_* expression with NH_4_^+^, regardless of the presence of NifL, indicated that there is another regulatory tier governing P*_nifLA_*. In various diazotrophic bacteria, *nifLA* expression is induced by a two-component system such as FixLJK, RegSR (RegBA/PrrAB), or NtrBC, with the latter expected for γ-proteobacteria [20]. Indeed, using BLASTp [46] to search for FixLJK (Accession: P23222-1, P29286) and RegSR (Accession: BAC46170-69) from *Bradyrhizobium diazoefficiens* USDA 110, and NtrBC from *K. pneumoniae* (Accession: CDO16416-7), we only identified homologs to NtrBC in *V. natriegens* (Table S1).

Across diverse bacteria, NtrBC governs a large regulon of nitrogen scavenging genes. In diazotrophs like *K. pneumoniae*, *P. stutzeri*, and *V. diazotrophicus*, this regulon includes *nif* genes (Fig 1A) [17, 20, 41, 47, 48], and deletion of *ntrBC* can dampen *nif* gene expression by an order of magnitude [17, 47]. We hypothesized that NtrBC controls *nifLA* in *V. natriegens* in response to NH_4_^+^ availability. To test this hypothesis, we deleted *ntrBC* and examined the growth of the resulting Δ*ntrBC* mutant with NH_4_^+^ versus N_2_. The Δ*ntrBC* mutant grew like the parent when provided NH_4_^+^ (Fig 4A), but a ∼16 h lag phase occurred under N_2_-fixing conditions (Fig 4B), after which the Δ*ntrBC* mutant exhibited similar growth trends as the parent (Fig 4B, C). We verified that the Δ*ntrBC* mutant growth was still dependent on NifA, as a Δ*ntrBC*Δ*nifA* strain was incapable of growth with N_2_ (Fig 4B).

**Fig 4.**
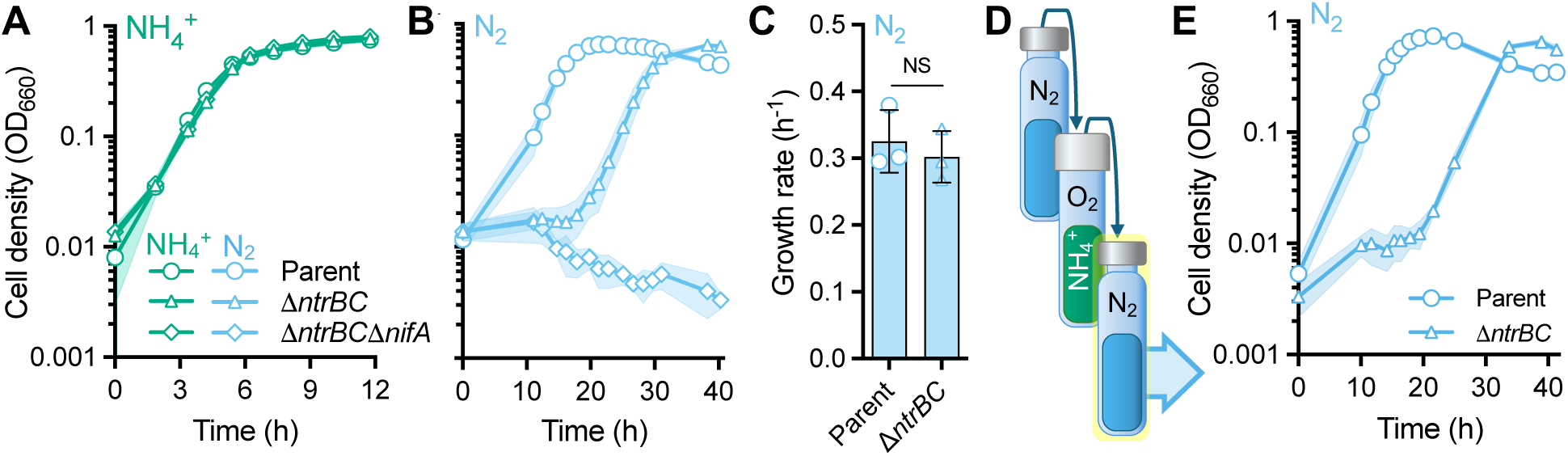
NtrBC is not required for diazotrophic growth. Growth curves for *V. natriegens* strains grown in anoxic minimal media with glucose and either NH_4_^+^ **(A)** or N_2_ **(B)**. Parent, NWH003 (Δ*dns*::Km^R^); Δ*ntrBC*, NWH014 (Δ*dns*::Km^R^); Δ*ntrBC*Δ*nifA*, NWH043 (Δ*dns*::Km^R^). **(A, B)** The same parent growth curves were used as in Fig 2. **(C)** Growth rates of the parent and Δ*ntrBC* mutant from panel B. NS, non-significant difference as determined using an unpaired, two-tailed t test. **(D)** Once N_2_-fixing cultures reached stationary phase, cells were subcultured into oxic media with NH_4_^+^, grown to stationary phase, then subcultured once more into anoxic minimal media with N_2_, where growth was then monitored as shown in panel **E**. **(A, B, E)** Points, mean; shading, SD; n=3.

To explain the lag phase and high diazotrophic growth rate of the Δ*ntrBC* mutant, we considered two possibilities: (i) suppressor mutations or (ii) *nif* gene expression is eventually activated by another protein(s) without mutation. We first tested the prediction that suppressor mutants, if present, would be enriched during growth with N_2_ and thus eliminate the Δ*ntrBC* lag phase upon subculturing. We thus grew Δ*ntrBC* cultures with N_2_ and then subcultured into aerobic conditions with NH_4_^+^ to produce progeny that were free of nitrogenase (Fig 4D); O_2_ irreversibly damages nitrogenase and NH_4_^+^ represses *nif* transcription [15, 20]. We then subcultured the aerobically grown cultures back to N_2_-fixing conditions (Fig 4D), where a similar lag phase was observed (Fig 4E), suggesting that the eventual Δ*ntrBC* mutant growth was not due to suppressor mutations. We thus suspected that another protein(s) compensated for the loss of *ntrBC* without mutation.

### The NifA regulon is a subset of the NtrC regulon

A possible candidate that could compensate for the absence of NtrBC is NifA, which can positively regulate its own expression in some diazotrophs, including *K. pneumoniae* [41–43, 49–51]. We thus sought to define the NtrBC and NifA regulons by using RNA-seq. To circumvent complications with the Δ*ntrBC* mutant lag phase and the inability of a Δ*nifA* mutant to grow with N_2_, we performed the analysis on non-growing cell suspensions, by transferring NH_4_^+^-grown cultures to nitrogen-free media under argon (Ar) to induce nitrogenase expression (Fig 5A). We verified that nitrogenase was expressed within 2 h by measuring H_2_ (Eq 1).

**Fig 5.**
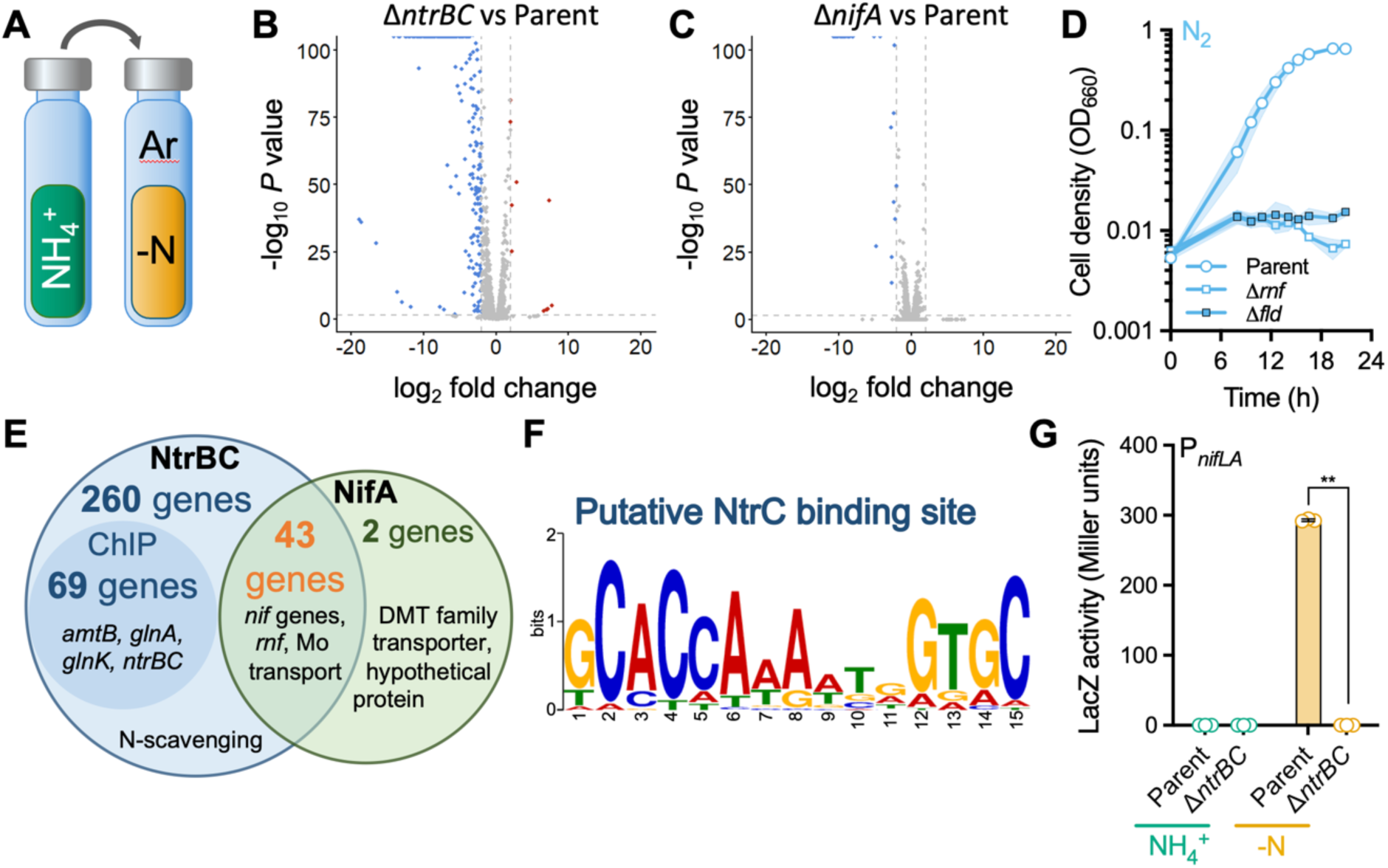
The NifA regulon is within the NtrBC regulon. **(A)** Cultures were grown anaerobically with glucose and NH_4_^+^, then were washed and resuspended in nitrogen-free media under argon to induce nitrogen starvation. **(B, C)** Differential gene expression analysis was performed on nitrogen-starved cell suspensions, comparing the Δ*ntrBC* mutant (NWH014, Δ*dns*::Km^R^) **(B)** or the Δ*nifA* mutant (NWH036, Δ*dns*::Km^R^) **(C)** to the parent (NWH003, Δ*dns*::Km^R^). Blue and red dots respectively indicate lower (log_2_ fold change ≤ -2) or higher (log_2_ fold change ≥ 2) mutant transcript levels (*P* ≤ 0.05). **(D)** Growth curves testing the essentiality of nitrogenase electron-transfer components encoded by the *rnf* operon and *nifF* in an anoxic minimal medium with N_2_ as the sole nitrogen source. Δ*rnf*, NWH038 (Δ*dns*::Km^R^); Δ*nifF*, NWH039 (Δ*dns*::Km^R^). Points, mean; shading, SD; n=3. **(E)** Comparison of the number of genes regulated by NtrBC and NifA, with some examples listed. The inner circle shows the number of genes that showd upstream binding by His-tagged NtrC as determined by ChIP-seq. **(F)** Putative NtrC DNA binding sequence based on binding sites from ChIP-seq. **(G)** A P*_nifLA_*-*lacZ* transcriptional reporter was used to measure *nifLA* expression in strains grown in a minimal anoxic medium with NH_4_^+^ or as non-growing cell suspensions (-N) after 2 h without nitrogen. Points, biological replicates; bars, mean; error bars, SD; n=3. The statistical difference was determined using an unpaired, two-tailed t test; **, *P* < 0.01.

Comparing the Δ*ntrBC* mutant and parent transcriptomes revealed 303 differentially expressed genes (DEGs) (Fig 5B, Table S2, File S1), 291 of which were downregulated in the Δ*ntrBC* mutant. These DEGs encompassed genes required for nitrogenase activity, including *nifA*, and for accessing other nitrogen sources. When comparing the Δ*nifA* mutant to the parent, a smaller regulon of 45 DEGs was observed, all of which were downregulated in the mutant (Fig 5C, Table S3, File S2). The NifA regulon consisted of all *nif* genes, and other genes needed for nitrogenase activity, such as those for an ABC molybdate transporter, an *rnf*-like electron transport complex, and the NifF flavodoxin; we verified that *rnf* and *nifF* are essential for N_2_ fixation using deletion mutants (Fig 5D). All but two of the NifA-dependent DEGs were within the NtrBC regulon (Fig 5E), indicating that NtrBC regulates N_2_ fixation via NifA.

The NtrBC regulon defined by RNA-seq contained other regulatory proteins. Thus, we sought to understand which genes are regulated through direct DNA-binding by NtrC. To identify NtrC binding sites, we ectopically expressed His-tagged NtrC in a Δ*ntrBC* mutant and performed chromatin immunoprecipitation combined with massively parallel sequencing (ChIP-seq) for cultures grown with N_2_ versus NH_4_^+^ (Fig S2). NtrC bound sites upstream of 69 genes /operons. All these genes were part of the NtrBC regulon defined by RNA-seq (Fig 5E, Table S2), and included *glnA*, encoding glutamine synthase; genes for the uptake and catabolism of alternative nitrogen sources like amino acids and purines; *glnK,* encoding a P_II_ protein; and *amtB1/amtB2*, encoding two NH_4_^+^ uptake transporters. The binding sites were common between N_2_-fixing conditions, where NtrC is presumably phosphorylated (Fig S2A, B), and conditions with NH_4_^+^, where NtrC is presumably unphosphorylated (Fig S2C, D). This observation is consistent with that from *Salmonella typhimurium* where DNA-binding by NtrC is independent of its phosphorylation state [52]. A putative *V. natriegens* NtrC DNA-binding motif was also identified (Fig 5F) that is present at least once in all NtrC-enriched regions and is similar to that in other bacteria [53, 54].

Despite NifA being within the NtrBC regulon according to RNA-seq, no NtrC binding was detected upstream of *nifLA* by ChIP-seq, nor was a binding motif identified upstream of *nifLA* or any of the other genes in the NifA regulon. To help resolve these conflicting results, we verified that *nifLA* expression was NtrC-dependent by measuring P*_nifLA_-lacZ* reporter activity. No P*_nifLA_-lacZ* activity was observed from the parent or the Δ*ntrBC* mutant during anoxic growth with NH_4_^+^, as expected. However, in non-growing cell suspensions that had been incubated without any nitrogen for 2 h, reporter activity was observed in the parent strain, but not in the Δ*ntrBC* mutant (Fig 5G). Thus, although ChIP-enrichment of *P_nifLA_* was not observed, our LacZ reporter data indicates that NtrBC is an activator of *nifLA* expression, at least within 2 h after transitioning to nitrogen-free conditions. Nonetheless, the dispensability of NtrBC for diazotrophic growth and the lack of binding sites suggests that it is not the only P*_nifLA_* activator.

### NifA can activate the expression of its own gene

Evidence from diazotrophs like *K. pneumoniae* [41–43] and higher P*_nifLA_*-*lacZ* activity in Δ*nifL* mutants (Fig 3) led us to hypothesize that NifA can activate *nifLA* expression (autoactivation). To test this hypothesis, we first examined the impact of NtrBC and NifA on P*_nifLA_*-*lacZ* reporter activity in non-growing cell suspensions after 2 h without nitrogen. As before, NtrBC was required for P*_nifLA_* expression within this time frame (Fig 5G and 6A). For the Δ*nifA* mutant, LacZ activity was still observed, but at 29% of parent levels. No reporter activity was detected from the Δ*ntrBC*Δ*nifA* mutant (Fig 6A). These results indicate that both NtrBC and NifA contribute to *nifLA* expression, but that NtrBC is required for expression within the first 2 h of nitrogen deprivation.

**Fig 6.**
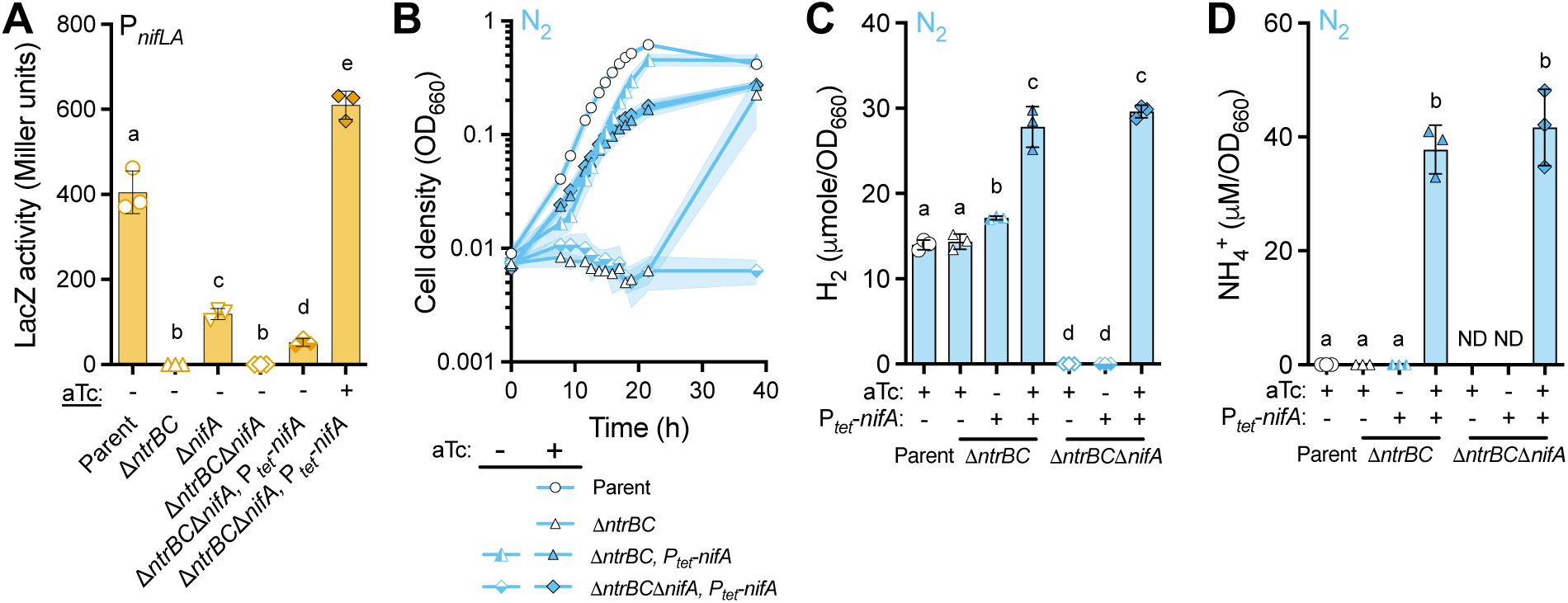
NifA activates its own expression. **(A)** A P*_nifLA_*-*lacZ* transcriptional reporter was used to measure *nifLA* expression in strains as nitrogen-starved cell suspensions with or without NifA expression from inducible P*_tet_*-*nifA*. Parent, NWH010 (Δ*dns*::Sp^R^*-*P*_nifLA_*-*lacZ*); Δ*ntrBC*, NWH015 (Δ*dns*::Sp^R^*-*P*_nifLA_*-*lacZ*); Δ*nifA*, NWH090 (Δ*dns*::Sp^R^*-*P*_nifLA_*-*lacZ*); Δ*ntrBC*Δ*nifA*, NWH126 (Δ*dns*::Sp^R^*-*P*_nifLA_*-*lacZ*); Δ*ntrBC*Δ*nifA*, *P_tet_-nifA*, NWH127 (Δ*dns*::Sp^R^*-P_nifLA_-lacZ-P_tet_-nifA-P_lacI_-tetR*) **(B)** Growth curves for strains grown in anoxic minimal media with glucose and or N_2_. Points, mean; shading, SD; n=3. **(C, D)** Accumulation of H_2_ in the headspace (limit of detection ∼10 nmol) **(C)** and of NH_4_^+^ in the supernatant (limit of detection ∼6 µM) **(D)** in stationary-phase (27 h) using separate cultures from those in panel B. **(A-D)** NifA expression was induced with anhydrotetracycline (aTc, +) or was not induced by adding DMSO (-). **(B-D)** Parent, NWH004 (Δ*dns*::Sp^R^); Δ*ntrBC*, NWH014 (Δ*dns*::Km^R^); Δ*nifA*, NWH036 (Δ*dns*::Km^R^); Δ*ntrBC*Δ*nifA* strains: NWH043 (Δ*dns*::Km^R^) and NWH108 (Δ*dns*::Sp^R^*-*P*_tet_-nifA*-P*_lacI_*-*tetR*). **(A, C, D)** Points, biological replicates; bars, mean; error bars, SD; n=3. Different letters indicate statistical differences as determined using a one-way ANOVA, *P* < 0.05. ND, not determined due to no growth.

Based on the above results, we surmised that initial activation by NtrC at P*_nifLA_* would increase NifA levels for subsequent autoactivation. If so, then expressing NifA from an inducible promoter would bypass the early need for NtrBC and activate P*_nifLA_*-*lacZ* expression. We tested this prediction by ectopically expressing *nifA* from an inducible promoter (P*_tet_*) in Δ*ntrBC*Δ*nifA* mutant cell suspensions that had been incubated without nitrogen for 2 h. Reporter activity was 1.5-times that of parent levels (Fig 6A). A small amount of P*_nifLA_*-*lacZ* activity was also observed in uninduced cultures (Fig 6A), likely due to leaky P*_tet_*-*nifA* expression. We conclude that NifA is sufficient to activate expression at P*_nifLA_*, thus serving as an autoactivator.

We hypothesized that NifA autoactivation could explain the Δ*ntrBC* mutant’s eventual diazotrophic growth (Fig 4B). For example, the Δ*ntrBC* mutant would be deficient for *nifLA* expression, but leaky or stochastic *nifLA* expression over the lag phase could eventually result in enough NifA in some cells to create a positive feedback loop of autoactivation. The time needed for this subpopulation to reach detectable levels would also contribute to the lag phase. Thus, we predicted that ectopic NifA expression should lessen the Δ*ntrBC* mutant lag phase under N_2_-fixing conditions. Even without induction, the Δ*ntrBC* mutant lag phase was reduced to < 10 h from the presence of P*_tet_::nifA* in the chromosome (Fig 6B), likely due to leaky P*_tet_*-*nifA* expression triggering further expression at the native *nifLA* operon (Fig 6A). In agreement with this notion, the removal of chromosomal *nifA* (Δ*ntrBC*Δ*nifA* mutant) required P*_tet_*-*nifA* induction for growth with N_2_ (Fig 6B). Induction of P*_tet_*-*nifA* in the Δ*ntrBC* mutant further eroded the lag period but also resulted in a lower growth rate and final cell density relative to the parent (Fig 6B). We speculated that the ectopic *nifA* expression led to abnormal nitrogenase activity. Indeed, when examining nitrogenase products (Eq 1), H_2_ levels were 2-fold higher in strains where P*_tet_*-*nifA* was induced compared to the parent (Fig 6C), and extracellular NH_4_^+^ was also observed (Fig 6D). Together, our results support a model where NtrC acts as an initial activator of *nifLA* expression in low nitrogen conditions, and then once expressed, NifA can further activate *nifLA* expression.

### PII proteins redundantly and negatively regulate nitrogenase

The above observations indicate that NtrBC makes a non-essential, but important, contribution to *V. natriegens* diazotrophic growth by initiating or amplifying early *nifLA* expression. However, it remained unclear what factors repress nitrogenase in response to NH_4_^+^. P_II_ proteins are often involved in sensing and transmitting fixed nitrogen availability. In *K. pneumoniae*, GlnB regulates nitrogenase expression by mediating the phosphorylation activity of NtrB, and GlnK alleviates NifL repression of NifA [20] (Fig 1A). We therefore investigated the contribution of *V. natriegens* P_II_ proteins to nitrogenase regulation.

We first tested whether *V. natriegens* P_II_ proteins affect growth with N_2_ versus NH_4_^+^ using single *glnB* or *glnK* deletion mutants. Both Δ*glnB* and Δ*glnK* mutants, grew like the parent, regardless of the nitrogen source (Fig 7A, B). Thus, single deletions did not cause a regulatory disruption, at least that was evident from growth trends. In contrast, a ΔPII mutant lacking both P_II_ proteins (Δ*glnB*Δ*glnK*), grew poorly with each nitrogen source (Fig 7C, D). Growth trends varied between biological replicates, suggesting that there was selective pressure for suppressor mutations.

**Fig 7.**
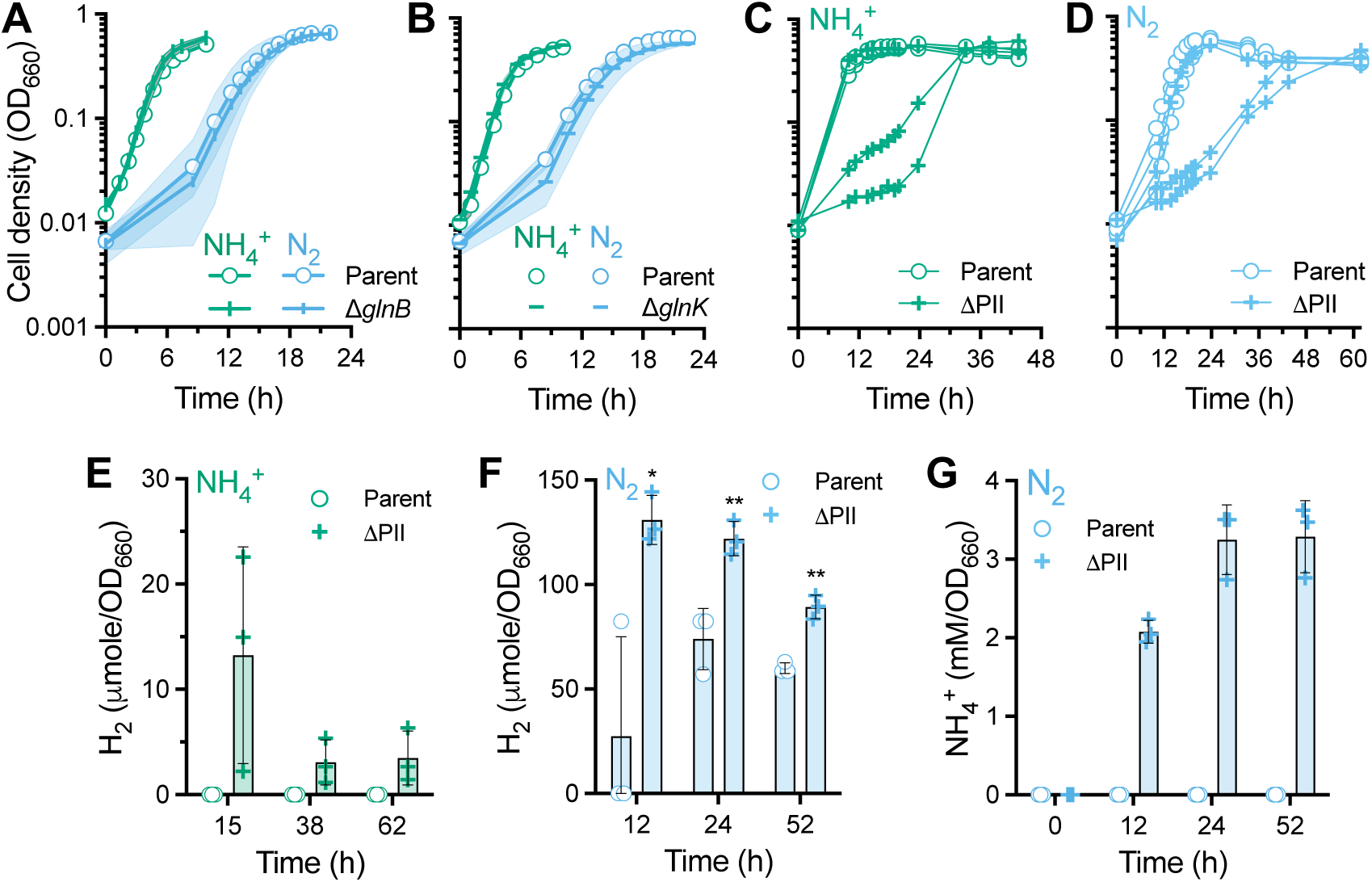
P_II_ proteins are redundant negative regulators of nitrogenase. (A-D) Growth curves for strains grown in anoxic minimal media with glucose and either NH_4_^+^ or N_2_. **(A, B)** Parent, NWH003 (Δ*dns*::Km^R^); Δ*glnB*, NWH042 (Δ*dns*::Km^R^), Δ*glnK*, NWH047 (Δ*dns*::Km^R^). Points, mean; shading, SD; n=3. **(C, D)** Parent, NWH004 (Δ*dns*::Sp^R^); ΔPII mutant, NWH050 (Δ*glnB*Δ*glnK*, Δ*dns*::Sp^R^). All replicate cultures are shown to reveal the diverse growth trends exhibited by the ΔPII mutant. **(E, F)** H_2_ was measured as an indicator of nitrogenase activity at different times for those cultures in panel C grown with NH_4_^+^ **(E)** or those in panel D grown with N_2_ **(F)**. **(G)** NH_4_^+^ in supernatant samples taken from N_2_-fixing cultures at different times. **(E-G)** Points, biological replicates; bars, mean; error bars, SD; n=3. Statistical differences were determined between the parent and the ΔPII mutant at each time point using an unpaired, two-tailed t test; *, *P* < 0.05; **, *P* < 0.01.

We hypothesized that the poor ΔPII mutant growth was due to overactive nitrogenase activity. Indeed, the ΔPII mutant produced H_2_ during growth with NH_4_^+^ (Fig 7E). H_2_ per cell was highest early in the growth curve, again supporting the notion that suppressor mutants with less nitrogenase activity were enriched during the experiment. Under N_2_-fixing conditions, ΔPII mutant H_2_ production was significantly higher than that of the parent (Fig 7F) and NH_4_^+^ was detected in the supernatant, accumulating up to 3.6 mM/OD_660_ (Fig 7G). These results indicate that *V. natriegens* P_II_ proteins have functional redundancy as negative regulators of nitrogenase.

P_II_ proteins can potentially affect broader aspects of nitrogen metabolism, for example by influencing the NtrBC-regulon. Thus, it was unclear if the ΔPII mutant growth defect was solely due to excessive nitrogenase activity or if other aspects of nitrogen metabolism were involved. To distinguish between these possibilities, we deleted either *nifA* or *ntrBC* in the ΔPII mutant and examined growth and nitrogenase activity via H_2_ production. When grown with NH_4_^+^, the ΔPIIΔ*nifA* mutant displayed a similar growth defect as the ΔPII mutant (Fig 8A) but it did not produce H_2_ (Fig 8B). This growth defect with NH_4_^+^ was eliminated in the ΔPIIΔ*ntrBC* mutant; growth and H_2_ production resembled the parent (Fig 8A, B). Similarly, in N_2_-fixing conditions, ΔPIIΔ*ntrBC* mutant growth and H_2_ production resembled that of the Δ*ntrBC* mutant rather than the ΔPII mutant, including a ∼16 h lag phase (Fig 8C, D). Thus, the ΔPII mutant growth defect is likely due to NtrBC contributing both to excessive nitrogenase and other aspects of nitrogen metabolism, which could include burdensome overexpression of nitrogen scavenging genes or dysregulation of assimilatory pathways.

**Fig 8.**
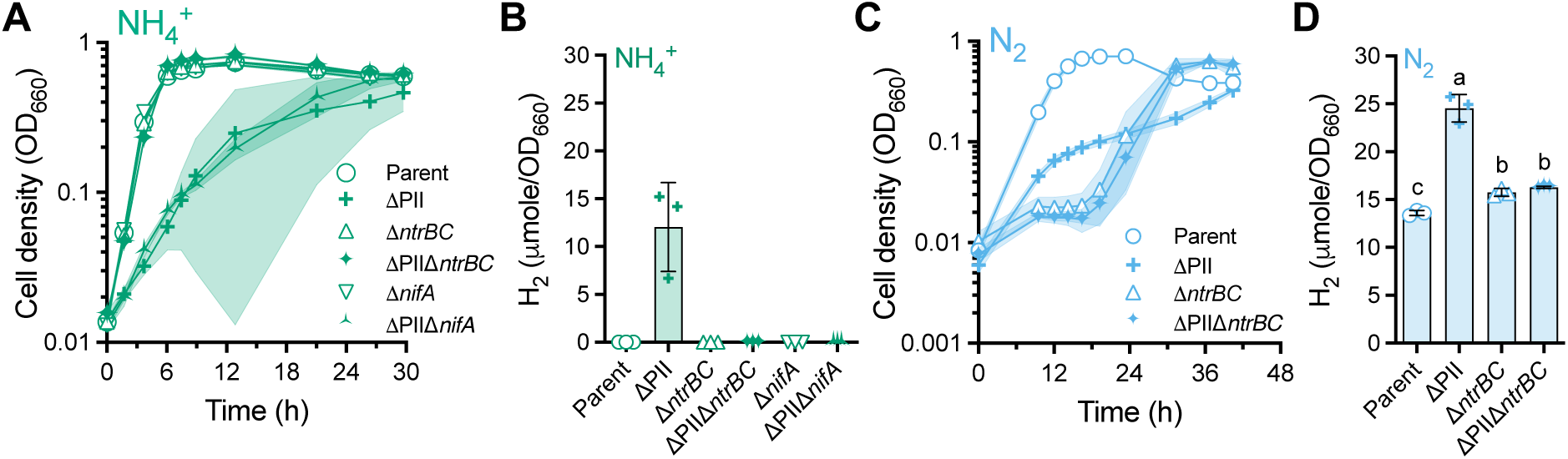
ΔPII growth defects are NtrBC-dependent. Growth curves **(A, C)** and final time-point H_2_ measurements **(B, D)** for strains grown in anoxic minimal media with glucose and either NH_4_^+^ **(A, B)** or N_2_ **(C, D)**. In N_2_ conditions, strains lacking *nifA* were omitted due to an inability to grow. Parent, NWH004 (Δ*dns*::Sp^R^); ΔPII, NWH050 (Δ*glnB*Δ*glnK*, Δ*dns*::Sp^R^); Δ*ntrBC*, NWH014 (Δ*dns*::Km^R^); ΔPIIΔ*ntrBC*, NWH111 (Δ*dns*::Sp^R^); ΔPIIΔ*nifA*, NWH112 (Δ*dns*::Sp^R^). **(A, C)** Points, mean; shading, SD; n=3. **(B, D)** Points, biological replicates; bars, mean; error bars, SD; n=3. **(D)** Different letters indicate statistical differences as determined using a one-way ANOVA, *P* < 0.05.

## DISCUSSION

### *V. natriegens* NifA autoactivation in comparison to other diazotrophs

We defined the basic architecture of nitrogenase regulation in *V. natriegens*. In some ways, the regulatory network resembles that of other diazotrophic γ-proteobacteria (Fig 1A and Fig 9). NtrBC facilitates the transition to N_2_-fixing conditions by responding to low nitrogen by activating early *nifLA* expression (Fig 6A and Fig 9). Without NtrBC, *nifA* can still be expressed by NifA autoactivation, but growth is delayed by ∼16 h (Fig 4B). Although we have not ruled out the possibility of an intermediate regulator participating between NtrBC and NifA, this would represent a highly unusual deviation from every nitrogenase regulatory scheme involving NtrBC. We thus find NifA autoactivation alone to be more likely. Additionally, P_II_ proteins were important negative regulators of nitrogenase expression that ensured that nitrogenase activity was responsive to NH_4_^+^ availability (Fig 8, 9).

**Fig 9.**
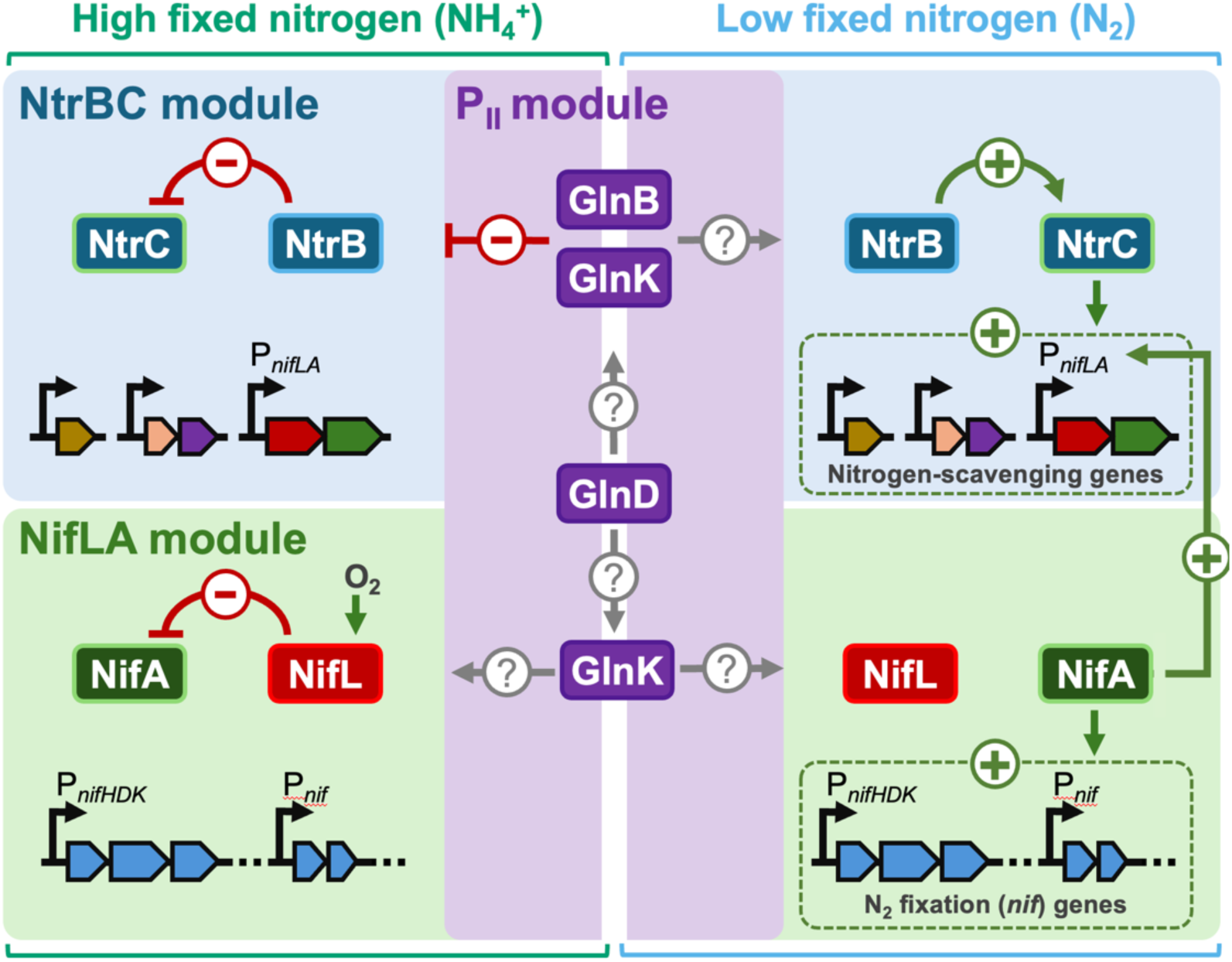
Emerging picture of nitrogenase regulation in *V. natriegens*. The NtrBC, NifLA, and P_II_ regulatory modules connect in ways that differ from other γ-proteobacteria. NtrBC is important for early *nif* gene expression but otherwise NifA autoactivation is sufficient for normal diazotrophic growth. P_II_ proteins play redundant negative roles, preventing NtrBC from contributing to excessive nitrogenase activity and broader growth defects. Several questions remain regarding how P_II_ proteins are regulated, including the role of post-translational modification by GlnD, and how else they might interact with the NtrBC and NifLA modules.

Although our work supports several findings from other diazotrophic γ-proteobacteria, it also highlights the importance of testing growth phenotypes. Early work on nitrogenase regulation, on which the field still heavily relies, often did not involve growth assays. For example, in *K. pneumoniae*, researchers described NifA-autoactivation from LacZ-reporter assays [42, 43] and from weak in-vitro NtrC binding to P*_nifLA_* relative to another NtrC-activated promoter, P*_glnA_* [41]. However, without growth phenotypes, one cannot truly know the relative contributions of regulators to nitrogenase activity. For example, in an α-proteobacterium where NifA autoactivation was suspected [30], an Δ*ntrBC* mutant had an order-of-magnitude lower diazotrophic growth rate than the parent [55]. We recommend that mutants for other model diazotrophs be revisited where growth phenotypes are lacking. For example, non-growing cell suspensions of a *P. stutzeri ntrB* mutant showed nitrogenase activity only after 20 h, similar to the lag time we observed in the *V. natriegens* Δ*ntrBC* mutant, and perhaps indicative of NifA autoactivation [47]. However, it is unknown if this trend translated to growth or if full nitrogenase activity was eventually achieved [47].

### *V. natriegens* nitrogenase regulation differs from other γ-proteobacteria

Although the *V. natriegens* nitrogenase regulatory network resembles that in other γ-proteobacteria, the contributions of the regulatory proteins differed. One example is the relatively minor impact of *V. natriegens* Δ*ntrBC* mutant aside from a diazotrophic lag phase (Fig 4B). Although diazotrophy is not yet well-characterized in *Vibrios*, a *V. diazotrophicus* Δ*ntrC* mutant exhibited 10% of the wild-type nitrogenase activity [17], a similar decrease as the α-proteobacterial mutant above [55]. This low activity still supported growth, but unfortunately no quantitative growth metrics are available to make comparisons [17]. Regardless, the defects of these mutants contrast our results where a *V. natriegens* Δ*ntrBC* mutant had a wild-type diaztotrophic growth rate (Fig 4C) and H_2_ production (Fig 6C), which should likely require wild-type nitrogenase activity.

Another way that *V. natriegens* nitrogenase regulation differs from other diazotrophic γ-proteobacteria is through the roles of P_II_ proteins. The *V. natriegens* P_II_ protein inventory of GlnB and GlnK resembles that of *K. pneumoniae*; *Pseudomonas* and *Azotobacter,* other well-characterized diazotrophic γ-proteobacteria, only have GlnK [56]. In *K. pneumoniae*, GlnB and GlnK play functionally opposing roles in nitrogenase regulation, while acting at different levels. GlnB negatively regulates nitrogenase by inhibiting NtrBC phosphorylation in response to fixed nitrogen (Fig 1); with fixed-nitrogen, a *K. pneumoniae glnB* mutant showed high P*_glnK_*-*lacZ* reporter activity, which is controlled by NtrC [28]. On the other hand, *K. pneumoniae* GlnK positively regulates nitrogenase by facilitating the dissociation of NifL from NifA (Fig 1) [26, 28]; for a *K. pneumoniae glnK* mutant, NifH-LacZ reporter activity was ∼25% of wild-type [28], even lower for a NifK-LacZ reporter [27], and diazotrophic growth was severely impaired [27]. *K. pneumoniae* GlnB also cannot substitute for GlnK, at least at physiological expression levels [25].

The above *K. pneumoniae* observations contrast those observed herein with *V. natriegens* where either P_II_ protein could be deleted with little consequence for diazotrophic growth, but deletion of both P_II_ proteins resulted in overactive nitrogenase activity that was dependent on NtrBC (Fig 7, 8). Thus, differing from the *K. pneumoniae* model, our results suggest that (i) each *V. natriegens* P_II_ protein plays a redundant negative role in regulating NtrBC and (ii) neither P_II_ protein is needed for dissociation of NifL from NifA. In these ways, *V. natriegens* P_II_ functions resemble those in the non-diazotrophic γ-proteobacterium *E. coli,* where GlnB and GlnK can play redundant regulatory roles and only disruption of both P_II_ proteins led to a NtrBC-dependent growth defect [57]. We also note that our ΔPII mutant resembles the α-proteobacterium *Rhodobacter capsulatus*, where deletion of both P_II_ proteins led to constitutive nitrogenase activity [33]. However, in *R. capsulatus* P_II_ protein regulation intersects with post-translational regulation of nitrogenase via DraTG, which *V. natriegens* ATCC 14048 does not possess. These differences from the expected *K. pneumoniae* model for nitrogenase regulation (Fig 1A) highlight the importance of characterizing nitrogenase regulation in diazotrophs of emerging interest. For *V. natriegens*, there is still much to understand about how the P_II_ proteins intersect with the other regulatory modules, including the effects of P_II_ modification with UMP via GlnD (Fig 9).

### Applied avenues for NH_4_^+^ production

Our work also identified P_II_ proteins as targets for engineering NH_4_^+^-excreting strains of *V. natriegens*. The highest level of NH_4_^+^ excretion was observed for the ΔP_II_ mutant, where extracellular NH_4_^+^ was ∼75-times higher than an Δ*ntrBC* mutant overexpressing NifA (Fig 7G vs Fig 6D). However, we suspect that the high nitrogenase activity, combined with dysregulation of broader nitrogen metabolism (Fig 8), made the desirable phenotype unstable; the poor growth created selective pressure for less NH_4_^+^ excretion. High NH_4_^+^ excretion is known to be unstable in other engineered NH_4_^+^-excreting bacteria [58–60]. Thus, further optimization of the trade-off between *V. natriegens* growth rate and NH_4_^+^ excretion is required. We anticipate that this pursuit will benefit from an improved understanding of interactions between P_II_ proteins and NtrBC and NifLA, along with a wholistic consideration of how NtrBC affects broader aspects of *V. natriegens* nitrogen metabolism.

## MATERIALS and METHODS

### Growth and cell suspension conditions

*V. natriegens* strains stored as 25% glycerol frozen stocks were struck for isolation on 1.5% agar plates of lysogeny broth (Miller) supplemented with 20 g NaCl L^-1^ (LB3), and antibiotics when appropriate (µg mL^-1^): 100 carbenicillin, 100 kanamycin (Km), 250 spectinomycin (Sp). Antibiotics and 100 µM isopropyl β-D-1-thiogalactopuranoide (IPTG) were also added to LB3 agar or broth as appropriate during strain construction. All experiments used minimal VMDC175 medium, which is M9-derived coculture medium [61] plus 100 mM MOPS (pH 7) and an additional 175 mM NaCl. Anoxic conditions were established by aliquoting 10 mL of media into 27 mL anaerobic tubes and bubbling with N_2_ or Ar. Tubes were sealed with rubber stoppers and aluminum crimps, then autoclaved. The following components were added after autoclaving via syringe (final concentrations): 25 mM glucose, 10 mM NH_4_Cl, 1 mM MgSO_4_, and 0.1 mM CaCl_2_. NH_4_Cl was omitted for N_2_-fixing conditions. When appropriate, 200 nM anhydrotetracycline hydrochloride (aTc) was added from a 20 µM stock solution in 100% dimethyl sulfoxide (DMSO) to induce expression from *P_tet_*; control cultures received the same volume of DMSO only. Single colonies were used to inoculate starter cultures in VMDC175 with either 1 or 10 mM NH_4_Cl, depending on whether the starter culture would be used to inoculate test conditions with N_2_ or NH_4_^+^, respectively.

After starter cultures reached late-exponential phase (0.65-0.85 OD_660_), they were used to inoculate test conditions at an initial cell density of ∼0.01 OD_660_. For nitrogen-starved cell suspensions, cultures were first grown to exponential-phase (0.3-0.5 OD_660_) in VMDC175 with 1 mM NH_4_Cl, then the entire culture was pelleted by centrifugation and washed once in an equal volume of VMDC175. Cell pellets were then resuspended in 0.3 mL of VMDC175 and the entire volume was transferred to 10 mL of anoxic VMDC175 under Ar with all media components except NH_4_Cl. All cultures and cell suspensions were incubated at 30°C, laid horizontally with shaking at 150 rpm for anoxic conditions or upright at a 45° slant with shaking at 250 rpm for oxic conditions. Assays involving cell suspensions were performed after 2 h, which we verified is adequate to consistently detect nitrogenase activity as H_2_. All shaking used a ¾” stroke length.

### Strain construction

Strains and primers are in Table S4 and Table S5, respectively. *V. natriegens* parent strains NWH003 or NWH004 were made by replacing *dns* (PN96_RS00885) in the type strain TND1964 with a kanamycin (Δ*dns*::Km^R^) or spectinomycin resistance cassette (Δ*dns*::Sp^R^), respectively. TND1964 is the “Dalia, SAD1302, 2016” variant wild-type *V. natriegens* ATCC 14048 with pMMB-*tfoX*, allowing for IPTG-inducible competency (https://portal.cultivarium.org/communities/vnat-sequencing?tab=data&dataset=Pilot%20study). Mutants were made as described [9] by natural transformation with both a deletion construct plus a selectable marker to replace the parent *dns* antibiotic cassette with the alternative cassette. Selectable markers were amplified from *V. natriegens* SAD1306 and were flanked by ∼1 kb upstream and downstream of *dns*. Deletion constructs were made by PCR amplifying ∼3 kb upstream and downstream of the region to be deleted using NEB Q5 DNA polymerase. The two regions were then connected to a 48 bp universal linker (MiniFRT; generated by combining oligos of each strand) by splicing-by-overlap extension (SOE) PCR using the outermost primers. The resulting SOE PCR product (1 µg) was co-transformed into the recipient strain along with 50 ng of either Δ*dns*::Sp^R^ or Δ*dns*::Km^R^. In all other cases, 200 ng of PCR-amplified DNA was transformed. To construct transcriptional reporters, *lacZ* plus 12 bp upstream of the start codon to capture the ribosomal binding site (RBS) was first amplified from pHRP309 [62]. Using SOE PCR, *lacZ* was then combined with an upstream Km^R^ cassette and flanked by ∼1 kb of homology surrounding *dns*. The product was then transformed into TND1964 generating NWH006 with a promoterless *lacZ* (Δ*dns*::Km^R^-*lacZ*). SOE PCR was then used to combine the Sp^R^ cassette upstream of a desired promoter, minus the native RBS, with 1 kb flanking DNA to match the region upstream of *dns* (upstream) and the *lacZ* insertion in NWH006 (downstream). The product was then transformed into NWH006 to generate a functional transcriptional reporter. To generate C-terminal tagged NtrC, *ntrBC* plus the native promoter and RBS (P*_ntrBC_*) was first amplified, fused with an upstream Km^R^ cassette and 1 kb flanking regions by SOE PCR as above to replace *dns* after transformation into TND1964, creating NWH060. The construct was then amplified and transformed into a Δ*ntrBC* mutant (NWH064). A PCR SOE product was also generated with a Sp^R^ cassette fused upstream of constitutive *glnA* promoter (P*_glnA_*) to replace the Km^R^-P*_ntrBC_* in NWH064, but preserving the native *ntrBC* RBS, creating NWH061. After verifying that the construct complemented diazotrophic lag phase, 1 kb upstream and downstream of the *ntrC* 3’ end were amplified and connected by a flexible Gly-Ser-Gly-Ser linker to a Hisx6 tag (generated by combining 30 bp oligos of each strand) directly upstream of the NtrC stop codon by SOE PCR. The final construct was then introduced into the Δ*ntrBC* mutant to create NWH071. To create an inducible *nifA* construct, a Sp^R^ cassette was combined upstream of an anhydrotetracycline-inducible *tet* promoter (P*_tet_*) and RBS amplified from pAJM.011 [63] and fused to *nifA* and a downstream constitutive P*_lacI_*-*tetR* repressor cassette (also from pAJM.011) by SOE PCR, creating a product with flanking regions to replace *dns*; pAJM.011 was a gift from C. Voigt (Addgene plasmid # 108529; http://n2t.net/addgene:108529 ; RRID:Addgene_108529). In all cases, transformants were isolated on LB3 agar with the appropriate antibiotic and verified by colony PCR and Sanger or Nanopore sequencing.

### β-galactosidase (LacZ) assays

Transcriptional LacZ reporter assays were performed in Z-buffer, consisting of 60 mM Na_2_HPO_4_, 40 mM NaH_2_PO_4_, 10 mM KCl, 1 mM MgSO_4_ pH adjusted to 7 with 1 M HCl before filter sterilization. Immediately prior to use, 2-mercaptoethanol was added to a final concentration of 50 mM. Lysis solution was prepared by diluting 10x FastBreak^TM^ Cell Lysis Reagent (Promega) in Z-Buffer with 10 mg mL^-1^ lysozyme. Samples from exponential-phase cultures or cell suspensions (both 0.3-0.5 OD_660_) were lysed by combining 20 µL of culture with 180 µL lysis solution in a 96-well plate. Lysis occurred over a 30 min incubation in a H1 Synergy microplate reader (BioTek) at 37°C with double-orbital shaking. A stock solution of o-nitrophenyl-β-D-galactopyranoside (ONPG) was prepared by dissolving ONPG in 100% DMSO, then diluting in Z-buffer to 4 mg mL^-1^. Following lysis, 20 µL ONPG solution was added to each well. LacZ activity was then tracked by absorbance (A_420_) for 1 h at 37°C in the plate reader. LacZ activity (V_420_ = A_420_ ÷ time) was quantified using the linear regression function in GraphPad Prism v.6 and expressed as Modified Miller units = (V_420_ • 1000 • 1.56) / (OD_660_ • 0.02 mL lysate volume) [64]. Values from appropriate control strains with a promoterless LacZ were subtracted from those from transcriptional reporters to control for ONPG instability.

### RNA extraction and sequencing

Cell suspensions were incubated under nitrogen-free conditions for 2 h at 30°C, then were chilled on ice and pelleted by centrifugation at 4°C. Supernatants were discarded and pellets frozen using dry ice before storing at -80°C. RNA was extracted with TRIzol^TM^ (Thermo Fisher) /chloroform as described [65]. Samples were treated with 4 U Turbo DNase (Invitrogen) in 100 µL at 37°C for 1 h, then purified using a QIAGEN RNeasy MinElute Cleanup Kit. RNA was quantified using an Agilent 2200 TapeStation at the Indiana University Center for Genomics and Bioinformatics (IU CGB), then submitted to SeqCoast Genomics where rRNA was depleted and libraries prepared using the Illumina Stranded Total RNA Prep Ligation Kit with Ribo-Zero Plus Microbiome and the RNA sequenced on an Illumina NextSeq2000 platform to produce 2×150 bp paired reads. Differential gene expression analysis was performed as described [66] using the *V. natriegens* ATCC 14048 genome (NCBI RefSeq: GCF_001456255.1). Differentially expressed genes with an adjusted *P*-value < 0.05 and a |log_2_(fold-change)| > 2.0 were considered significant.

### Chromatin immunoprecipitation-sequencing (ChIP-seq)

ChIP-seq was performed as described [67, 68] with some modifications. Five 10-mL anaerobic cultures were grown to exponential phase (0.3-0.5 OD_660_). Samples (2 ml) were then extracted and pooled for whole-genome sequencing to be used as the ‘input’ for ChIP-seq normalization. These pooled samples were pelleted by centrifugation at 4°C, supernatants discarded, pellets frozen in dry ice, and then stored at -80°C. gDNA was extracted using the QIAGEN DNeasy Blood and Tissue kit. The remaining anaerobic cultures received an injection of 37% formaldehyde (1% final) and were crosslinked for 30 min at room temperature. Tubes were then unsealed, cultures pooled, and crosslinking quenched with 125 mM glycine. Cells were then pelleted by as above, resuspended in 1 mL ice-cold TBS buffer (20 mM TRIS, 0.9% NaCl, pH 7.4), transferred to a 1.5 ml tube and pelleted again by centrifugation at 4 °C. This wash step was repeated, then cell pellets were frozen using dry ice, and stored at -80°C. Chromatin was immunoprecipitated from cell pellets as described [67], except Protein G Mag Sepharose^TM^ Xtra magnetic beads (Cytiva) and anti-His antibodies (Genscript) were used to pull down His-tagged NtrC. The strain NHW061, with untagged NtrC was used as a control for nonspecific enrichment. Library preparation for genomic and ChIP DNA was performed using the NEBNext Ultrall kit (NEB) and sequenced at the IU CGB using an Illumina NextSeq2000, generating paired-end reads. Reads were mapped to the *V. natriegens* ATCC 14048 genome (GCA_001456255.1) using CLC Genomics Workbench (CLC Bio, QIAGEN). ChIP and input sequences were normalized based on the total number of reads, and ChIP enrichment (ChIP/input) was plotted using customized R scripts (https://github.com/xindanwanglab/Haas-2026a).

### Analytical procedures

Cell density was measured as turbidity at 660 nm (OD_660_) using a Genesys 20 spectrophotometer (Thermo-Fisher) directly in culture tubes. Growth rates were calculated using GraphPad Prism v.6 by fitting an exponential trendline to turbidity measurements between 0.1-0.8 OD_660_. H_2_ was quantified by sampling 0.1 mL of headspace using a gas-tight syringe and injecting into a Shimadzu GC-2014 gas chromatograph with a thermal conductivity detector as described [69]. The detection limit was 10 nmol H_2_. NH_4_^+^ was quantified with an indophenol colorimetric assay [61] using late-stationary phase cultures (0.65-0.85 OD_660_) grown in VMDC175 but omitting MOPS and using 15 mM glucose to avoid acidic conditions that interfere with the assay. Culture samples (1 ml) were pelleted, and 550 µL of supernatant was combined with 50 µL of 1 M NaOH, 100 µL phenol nitroprusside (Sigma-Aldrich) and 100 µL of alkaline hypochlorite (Sigma-Aldrich). For ΔPII mutant samples only, 50 µL of supernatant plus 500 µL of VMDC175 (no MOPS) was used. Samples were vortexed and incubated at room temperature for 15 minutes, then absorbance measured (A_630_) in 1 mL cuvettes. Standard curves used NH_4_Cl in VMDC175 (no MOPS). The limit of detection was ∼6 µM NH_4_^+^.

### Statistics

GraphPad Prism v.6 was used for all statistical analyses for growth, LacZ-reporter, H_2_, and NH_4_^+^ assays.

## Data availability

Processed files and raw reads for RNA-seq and ChIP-seq have been deposited in NCBI’s Genome Expression Omnibus [70] with GEO series accession numbers GSE333724 and GSE333729, respectively. NCBI Sequence Read Archive and under BioProject accession number PRJNA1472683 (https://www.ncbi.nlm.nih.gov/bioproject).

## ACKNOWLEDGEMENTS

This work was supported in part by National Science Foundation grants MCB-1749489 (JBM; CAREER) and DBI-2022049 (XW; Biology Integration Institutes Program), National Institutes of Health grants R35GM128674 (ABD) and R01GM141242, R01GM143182, and R01AI172822 (XW), the US Army Research Office grant W911NF-17-1-0159 (JBM), an IU Kindig award (NWH), and the IU College of Arts and Sciences. Supercomputing resources were supported in part by Lilly Endowment, Inc., through its support for the IU Pervasive Technology Institute.

We are grateful to the IU CGB for WGS sequencing support, to J. Lewis for contributions to mutant construction, and to C. Fuqua, C. Landeta, the IU Biology *Vibrio* group, and the McKinlay lab for helpful discussions.

## Notes

### Competing Interest Statement

The authors have declared no competing interest.

