## Supplementary material for "Nitrogenase regulation in *Vibrio natriegens* differs from other γ-proteobacteria": Table S

Contents:

**Fig S1. NifL prevents nitrogenase expression in response to $O_2$**

**Fig S2. NtrC binds the same chromosomal (Ch) sites when *V. natriegens* is grown with $N_2$ versus $NH_4^+$**

**Table S1. *V. natriegens* genes expected to be involved in $N_2$ fixation**

**Table S2. Differentially expressed genes in nitrogen-starved Δ*ntrBC* mutant versus the parent**

**Table S3. Differentially expressed genes in nitrogen-starved Δ*nifA* mutant versus the parent**

**Table S4. Bacterial strains**

**Table S5. Primers**

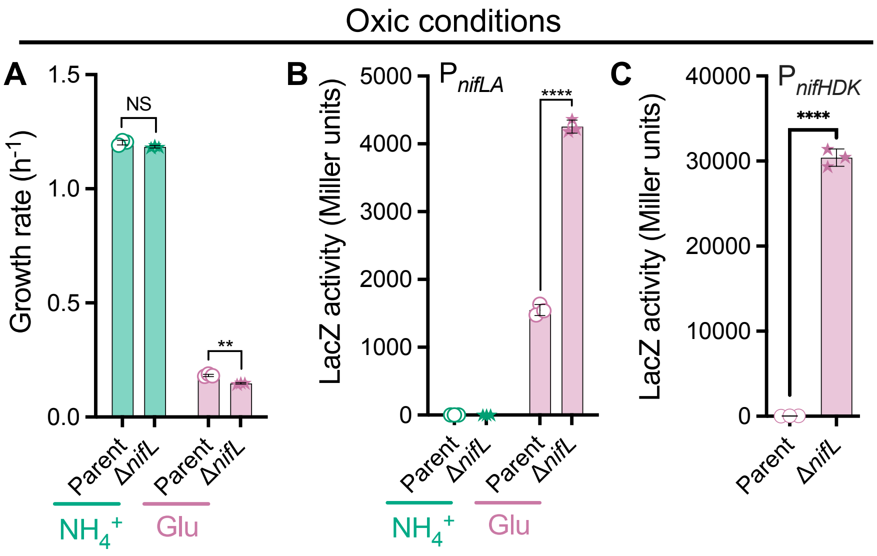


**Fig S1. NifL prevents nitrogenase expression in response to $O_2$. (A)** *V. natriegens* growth rates in oxic minimal media with glucose and either $NH_4^+$ or glutamate as the nitrogen source. Parent, NWH003 (Δ*dns*::$Km^R$); Δ*nifL*, NWH037 (Δ*dns*::$Km^R$). **(B, C)** A *lacZ* transcriptional reporter was used to compare expression from $P_{nifLA}$ **(B)** or $P_{nifHDK}$ **(C)**. **(A – C)** Points, biological replicates; bars, mean; error bars, SD; n=3. Statistical differences were determined using an unpaired, two-tailed t tests; NS, non-significant; **, *P* < 0.01; ****, *P* < 0.0001.

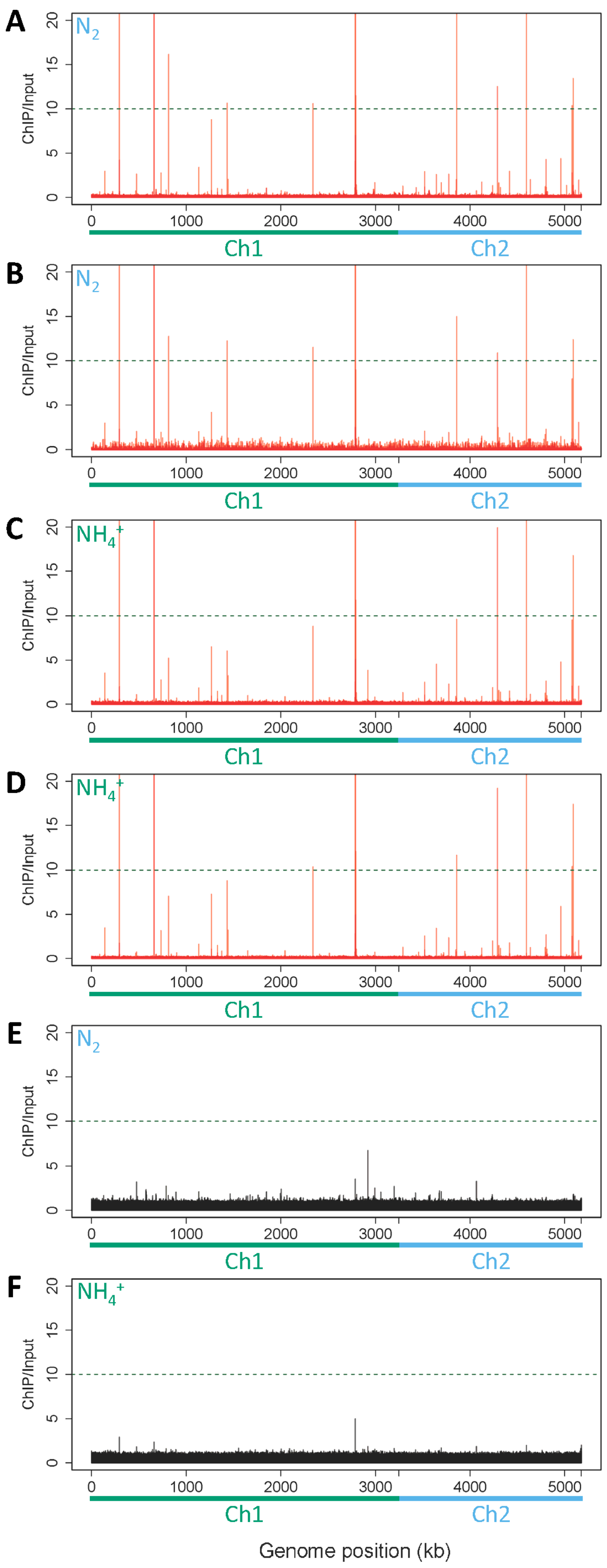


**Fig S2. NtrC binds the same chromosomal (Ch) sites when *V. natriegens* is grown with $N_2$ versus $NH_4^+$.** DNA binding of His-tagged NtrC was as determined by ChIP-seq using strain NWH071. Each plot with red bars shows a biological with $N_2$ **(A, B)** or $NH_4^+$ **(C, D)**. Plots with black bars are controls from strain NWH061 with untagged NtrC grown with $N_2$ **(E)** or $NH_4^+$ **(F)**. Sequencing coverage values were determined for 1 kb bins across the genome for ChIP-enriched samples, and the coverage ratio relative to a whole-genome control is plotted on the Y-axis.

**Table S1. *V. natriegens* genes expected to be involved in $N_2$ fixation**[a]

| Gene (synonyms) | Description | Locus tag | Protein ID |
|---|---|---|---|
| **Regulatory proteins** | | | |
| *rpoN* (*ntrA*, *glnF*) | σN/σ54 | PN96_RS00495 | WP_020334488 |
| *ntrB* (*glnL*, *glnR*) | Nitrogen regulatory protein NR(II) | PN96_RS12940 | WP_014230519 |
| *ntrC* (*glnG*, *glnT*) | Nitrogen regulatory protein NR(I) | PN96_RS12945 | WP_014230518 |
| *glnD* | Bifunctional uridylyltransferase/uridylyl-removing enzyme | PN96_RS02290 | WP_020335150 |
| *glnB* | Nitrogen regulatory protein P-II | PN96_RS02395 | WP_014232746 |
| *glnK* | P-II family nitrogen regulator | PN96_RS01485 | WP_005386570 |
| *nifA* | *nif*-specific transcriptional activator | PN96_RS19610 | WP_020333834 |
| *nifL* | NifA anti-activator | PN96_RS19615 | WP_020333835 |
| **Transporters** | | | |
| *amtB1* | Ammonium transporter | PN96_RS01490 | WP_014232922 |
| *amtB2* | Ammonium transporter | PN96_RS05910 | WP_020333273 |
| *modA1* | Molybdate ABC transporter, substrate-binding protein | PN96_RS19520 | WP_020333821 |
| *modB1* | Molybdate ABC transporter, permease subunit | PN96_RS19525 | WP_231119409 |
| *modC1* | Molybdate ABC transporter, ATP-binding protein | PN96_RS19530 | WP_020333823 |
| *modA2* | Molybdate ABC transporter, substrate-binding protein | PN96_RS20775 | WP_020334863 |
| *modB2* | Molybdate ABC transporter, permease subunit | PN96_RS20780 | WP_020334865 |
| *modC2* | Molybdate ABC transporter, ATP-binding protein | PN96_RS20785 | WP_024372990 |
| **Nitrogenase biosynthesis** | | | |
| *nifH* | Nitrogenase Fe protein | PN96_RS19440 | WP_020333808 |
| *nifD* | Nitrogenase Mo-Fe protein, α-subunit | PN96_RS19445 | WP_014234003 |
| *nifK* | Nitrogenase Mo-Fe protein, β-subunit | PN96_RS19450 | WP_014234004 |
| *nifT* | Nitrogen fixation protein | PN96_RS19455 | WP_020333810 |
| *RS19460* | NifB/NifX family Mo-Fe cluster-binding protein | PN96_RS19460 | WP_020333811 |
| *RS19465* | Hypothetical protein | PN96_RS19465 | WP_020333812 |
| *nifE* | Nitrogenase Mo-Fe cofactor biosynthesis protein | PN96_RS19470 | WP_020333813 |
| *nifN* | Nitrogenase Mo-Fe cofactor biosynthesis protein | PN96_RS19475 | WP_020333814 |
| *RS19480* | NifB/NifX family Mo-Fe cluster-binding protein | PN96_RS19480 | WP_014234009 |
| *nifU* | Fe-S cluster assembly protein | PN96_RS19490 | WP_020333815 |
| *nifS* | Cysteine desulfurase | PN96_RS19495 | WP_020333816 |
| *nifV* | Homocitrate synthase | PN96_RS19500 | WP_020333817 |
| *nifW* | Nitrogenase-stabilizing/protective protein | PN96_RS19505 | WP_020333818 |
| *nifZ* | Nitrogen fixation protein | PN96_RS19510 | WP_020333819 |
| *nifM* | Peptidylprolyl isomerase | PN96_RS19515 | WP_020333820 |
| *nifQ* | Nitrogenase Mo-Fe cofactor biosynthesis protein | PN96_RS19595 | WP_020333832 |
| *nifB* | Nitrogenase Mo-Fe cofactor biosynthesis protein | PN96_RS19605 | WP_236614651 |
| **Electron transfer** | | | |
| *fdxB* | Ferredoxin III, *nif*-specific | PN96_RS19485 | WP_014234010 |
| *RS24415* | Putative ferredoxin | PN96_RS24415 | WP_076633454 |
| *nifF* | Flavodoxin | PN96_RS19655 | WP_020333842 |
| *rnfA* | RnfABCDGE type electron transport complex, subunit A | PN96_RS19620 | WP_014234038 |
| *rnfB* | RnfABCDGE type electron transport complex, subunit B | PN96_RS19625 | WP_020333836 |
| *rnfC* | RnfABCDGE type electron transport complex, subunit C | PN96_RS19630 | WP_020333837 |
| *rnfD* | RnfABCDGE type electron transport complex, subunit D | PN96_RS19635 | WP_020333838 |
| *rnfG* | RnfABCDGE type electron transport complex, subunit G | PN96_RS19640 | WP_020333839 |
| *rnfE* | RnfABCDGE type electron transport complex, subunit E | PN96_RS19645 | WP_020333840 |
| *rnfH* | RnfH family protein | PN96_RS19650 | WP_020333841 |

[a]*V. natriegens* proteins were identified based on existing annotations or from manual BLASTp hits that had > 25% identity across > 50% of the query sequence (i.e., NtrBC).

**Table S2. Differentially expressed genes in nitrogen-starved Δ*ntrBC* mutant versus the parent[a]**

| Locus tag | Gene name | Description | $Log_2$ fold-change |
|---|---|---|---|
| **Downregulated genes in Δ*ntrBC*** | | | |
| **PN96_RS12940** | ***ntrB* (*glnL*)** | **nitrogen regulation protein NR(II)** | **-18.90** |
| **PN96_RS12945** | ***ntrC* (*glnG*)** | **nitrogen regulation protein NR(I)** | **-18.61** |
| PN96_RS19430 | *cobA* | uroporphyrinogen-III C-methyltransferase | -16.54 |
| PN96_RS17460 | *eutB* | ethanolamine ammonia-lyase subunit EutB | -14.08 |
| PN96_RS19625 | *rnfB* | RnfABCDGE type electron transport complex subunit B | -13.63 |
| PN96_RS17465 | *eutC* | ethanolamine ammonia-lyase subunit EutC | -13.41 |
| PN96_RS19440 | *nifH* | nitrogenase Fe protein | -13.30 |
| PN96_RS19635 | *rnfD* | RnfABCDGE type electron transport complex subunit D | -12.94 |
| PN96_RS19620 | *rnfA* | RnfABCDGE type electron transport complex subunit A | -12.85 |
| PN96_RS19630 | *rnfC* | RnfABCDGE type electron transport complex subunit C | -12.47 |
| PN96_RS19445 | *nifD* | nitrogenase Fe-Mo protein α-chain | -12.44 |
| PN96_RS19490 | *nifU* | Fe-S cluster assembly protein NifU | -12.37 |
| PN96_RS19450 | *nifK* | nitrogenase Fe-Mo protein β-subunit | -12.03 |
| PN96_RS19495 | *nifS* | cysteine desulfurase NifS | -11.93 |
| PN96_RS16310 | | PLP-dependent aminotransferase family protein | -11.93 |
| **PN96_RS17800** | | **nucleobase:cation symporter-2 family protein** | **-11.85** |
| PN96_RS16080 | | AraC family transcriptional regulator | -11.76 |
| PN96_RS19640 | *rnfG* | RnfABCDGE type electron transport complex subunit G | -11.52 |
| PN96_RS19455 | *nifT* | putative nitrogen fixation protein NifT | -11.44 |
| PN96_RS19500 | *nifV* | homocitrate synthase | -11.29 |
| PN96_RS19505 | *nifW* | nitrogenase-stabilizing/protective protein NifW | -11.28 |
| **PN96_RS06645** | | **helix-turn-helix domain-containing protein** | **-11.16** |
| PN96_RS19510 | *nifZ* | nitrogen fixation protein NifZ | -10.91 |
| PN96_RS19515 | *nifM* | peptidylprolyl isomerase | -10.88 |
| **PN96_RS20310** | | **transporter substrate-binding protein** | **-10.87** |
| PN96_RS19645 | *rnfE* | RnfABCDGE type electron transport complex subunit E | -10.74 |
| PN96_RS19605 | *nifB* | nitrogenase cofactor biosynthesis protein NifB | -10.67 |
| PN96_RS19615 | *nifL* | nitrogen fixation negative regulator NifL | -10.66 |
| PN96_RS19595 | *nifQ* | nitrogen fixation protein NifQ | -10.61 |
| PN96_RS17470 | *eat* | ethanolamine permease | -10.61 |
| PN96_RS19460 | | NifB/NifX family Fe-Mo cluster-binding protein | -10.61 |
| PN96_RS24200 | | hypothetical protein | -10.56 |
| PN96_RS19470 | *nifE* | nitrogenase Fe-Mo cofactor biosynthesis protein NifE | -10.56 |
| PN96_RS19650 | *rnfH* | RnfH family protein | -10.53 |
| PN96_RS19520 | *modA1* | molybdate ABC transporter substrate-binding protein | -10.51 |
| **PN96_RS03150** | | **PLP-dependent aspartate aminotransferase family protein** | **-10.44** |
| PN96_RS19525 | *modB1* | molybdate ABC transporter permease subunit | -10.42 |
| PN96_RS19465 | | hypothetical protein | -10.40 |
| PN96_RS19655 | *nifF* | flavodoxin | -10.25 |
| PN96_RS19530 | *modC1* | molybdenum ABC transporter ATP-binding protein | -10.02 |

| | | | |
|---|---|---|---|
| PN96_RS19475 | *nifN* | nitrogenase Fe-Mo cofactor biosynthesis protein NifN | -10.01 |
| **PN96_RS20315** | | **amino acid ABC transporter permease** | **-9.95** |
| **PN96_RS01485** | ***glnK*** | **PII family nitrogen regulator** | **-9.92** |
| PN96_RS19610 | *nifA* | nif-specific transcriptional activator NifA | -9.89 |
| PN96_RS19480 | | NifB/NifX family Fe-Mo cluster-binding protein | -9.83 |
| PN96_RS19485 | *fdxB* | ferredoxin III, nif-specific | -9.60 |
| **PN96_RS17830** | ***xdhA*** | **xanthine dehydrogenase small subunit** | **-9.58** |
| PN96_RS03145 | | transporter substrate-binding domain-containing protein | -9.58 |
| PN96_RS17455 | | helix-turn-helix domain-containing protein | -9.49 |
| PN96_RS03475 | | hypothetical protein | -9.36 |
| **PN96_RS19750** | | **AraC family transcriptional regulator** | **-9.26** |
| **PN96_RS20320** | | **amino acid ABC transporter permease** | **-9.22** |
| **PN96_RS17815** | | **2-oxo-4-hydroxy-4-carboxy-5-ureidoimidazoline decarboxylase** | **-9.03** |
| **PN96_RS17825** | ***xdhB*** | **xanthine dehydrogenase molybdopterin binding subunit** | **-8.98** |
| **PN96_RS17820** | ***xdhC*** | **xanthine dehydrogenase accessory protein XdhC** | **-8.86** |
| **PN96_RS17810** | ***uraH*** | **hydroxyisourate hydrolase** | **-8.86** |
| **PN96_RS03140** | | **amino acid ABC transporter permease** | **-8.79** |
| **PN96_RS01490** | ***amtB1*** | **ammonium transporter** | **-8.77** |
| PN96_RS03920 | | hypothetical protein | -8.71 |
| **PN96_RS17805** | | **urate hydroxylase PuuD** | **-8.60** |
| **PN96_RS20325** | | **amino acid ABC transporter ATP-binding protein** | **-8.38** |
| **PN96_RS10880** | | **ABC transporter substrate-binding protein** | **-8.13** |
| **PN96_RS03795** | | **circularly permuted type 2 ATP-grasp protein** | **-7.92** |
| **PN96_RS10885** | | **ABC transporter ATP-binding protein** | **-7.91** |
| PN96_RS18585 | | muconate/chloromuconate family cycloisomerase | -7.75 |
| **PN96_RS21135** | ***glgC*** | **glucose-1-phosphate adenylyltransferase** | **-7.70** |
| **PN96_RS03135** | | **amino acid ABC transporter ATP-binding protein** | **-7.56** |
| **PN96_RS03790** | | **α-E domain-containing protein** | **-7.56** |
| PN96_RS16315 | | TSUP family transporter | -7.48 |
| PN96_RS19425 | *nirB* | nitrite reductase large subunit NirB | -7.46 |
| **PN96_RS10890** | | **ABC transporter permease** | **-7.43** |
| **PN96_RS03785** | | **transglutaminase family protein** | **-7.26** |
| PN96_RS04685 | | glycosyltransferase | -7.25 |
| **PN96_RS23215** | ***gabT*** | **4-aminobutyrate--2-oxoglutarate transaminase** | **-7.23** |
| PN96_RS06690 | | AraC family transcriptional regulator | -7.20 |
| PN96_RS16070 | | RidA family protein | -7.19 |
| **PN96_RS19740** | | **hypothetical protein** | **-7.18** |
| PN96_RS19420 | *nirD* | nitrite reductase small subunit NirD | -7.15 |
| **PN96_RS23225** | | **TRAP transporter substrate-binding protein** | **-6.99** |
| **PN96_RS10875** | | **amidohydrolase family protein** | **-6.97** |
| PN96_RS23370 | | AraC family ligand binding domain-containing protein | -6.93 |
| **PN96_RS10895** | | **2-oxoglutarate and Fe-dependent oxygenase domain-containing protein** | **-6.89** |

| | | | |
|---|---|---|---|
| **PN96_RS23230** | | **TRAP transporter small permease** | **-6.72** |
| **PN96_RS19745** | ***putA*** | **bifunctional proline dehydrogenase/L-glutamate gamma-semialdehyde dehydrogenase PutA** | **-6.67** |
| PN96_RS19035 | *guaD* | guanine deaminase | -6.53 |
| **PN96_RS06405** | | **NCS1 family nucleobase:cation symporter-1** | **-6.47** |
| PN96_RS19415 | | NarK family nitrate/nitrite MFS transporter | -6.34 |
| PN96_RS05590 | | amino acid ABC transporter substrate-binding protein | -6.29 |
| **PN96_RS20305** | | **MurR/RpiR family transcriptional regulator** | **-6.26** |
| PN96_RS19435 | | nitrate reductase | -6.23 |
| PN96_RS16320 | | SDR family NAD(P)-dependent oxidoreductase | -6.20 |
| PN96_RS04760 | | STAS domain-containing protein | -6.20 |
| **PN96_RS17835** | | **TetR/AcrR family transcriptional regulator** | **-6.20** |
| **PN96_RS23210** | | **PLP-dependent aminotransferase family protein** | **-6.18** |
| **PN96_RS22690** | | **GlxA family transcriptional regulator** | **-6.17** |
| PN96_RS04680 | | glycosyltransferase family 2 protein | -6.11 |
| **PN96_RS23220** | | **NAD-dependent succinate-semialdehyde dehydrogenase** | **-5.98** |
| PN96_RS22045 | | phytanoyl-CoA dioxygenase family protein | -5.86 |
| **PN96_RS23235** | | **TRAP transporter large permease subunit** | **-5.84** |
| PN96_RS05585 | | amino acid ABC transporter permease | -5.80 |
| PN96_RS11180 | *ung* | uracil-DNA glycosylase | -5.72 |
| PN96_RS04700 | | acyltransferase | -5.70 |
| PN96_RS05580 | | amino acid ABC transporter permease | -5.63 |
| PN96_RS07685 | | hybrid-cluster NAD(P)-dependent oxidoreductase | -5.62 |
| PN96_RS04755 | | OmpA family protein | -5.58 |
| **PN96_RS19730** | ***aspA*** | **aspartate ammonia-lyase** | **-5.55** |
| PN96_RS22005 | | DUF4440 domain-containing protein | -5.50 |
| PN96_RS24300 | | hypothetical protein | -5.49 |
| **PN96_RS19735** | ***putP*** | **sodium/proline symporter PutP** | **-5.46** |
| PN96_RS05575 | | amino acid ABC transporter ATP-binding protein | -5.40 |
| PN96_RS04715 | | glycosyltransferase family 4 protein | -5.30 |
| **PN96_RS12935** | | **DUF4124 domain-containing protein** | **-5.16** |
| **PN96_RS23265** | ***dgcN*** | **N-acetyltransferase DgcN** | **-5.15** |
| PN96_RS07680 | *hcp* | hydroxylamine reductase | -5.09 |
| PN96_RS21995 | | hypothetical protein | -5.08 |
| **PN96_RS21975** | | **FAD-dependent oxidoreductase** | **-5.07** |
| PN96_RS15975 | | aromatic amino acid transporter | -5.06 |
| PN96_RS22010 | | SDR family oxidoreductase | -4.91 |
| PN96_RS15970 | *tnaA* | tryptophanase | -4.90 |
| **PN96_RS23270** | ***dgcA*** | **N-acetyl-D-Glu racemase DgcA** | **-4.90** |
| PN96_RS19410 | | bifunctional protein-serine/threonine kinase/phosphatase | -4.86 |
| PN96_RS22015 | | S-(hydroxymethyl)glutathione dehydrogenase/class III alcohol dehydrogenase | -4.84 |
| PN96_RS04710 | | oligosaccharide flippase family protein | -4.73 |
| **PN96_RS23275** | | **D-amino-acid transaminase** | **-4.73** |

| | | | |
|---|---|---|---|
| **PN96_RS01495** | | **$Fe^{3+}$ ABC transporter substrate-binding protein** | **-4.72** |
| PN96_RS04720 | | glycosyltransferase family 4 protein | -4.72 |
| PN96_RS04675 | | sugar transferase | -4.71 |
| **PN96_RS21970** | | **flavodoxin domain-containing protein** | **-4.67** |
| **PN96_RS05910** | ***amtB2*** | **ammonium transporter** | **-4.67** |
| PN96_RS06400 | | amidase | -4.60 |
| **PN96_RS06410** | | **GntR family transcriptional regulator** | **-4.48** |
| PN96_RS13155 | | Flp family type IVb pilin | -4.41 |
| PN96_RS23060 | | helix-turn-helix domain-containing protein | -4.28 |
| PN96_RS19360 | | choline ABC transporter substrate-binding protein | -4.25 |
| PN96_RS20515 | *dld* | D-lactate dehydrogenase | -4.15 |
| PN96_RS22030 | | cytochrome P450 | -4.14 |
| **PN96_RS06395** | | **gamma-glutamyltransferase family protein** | **-4.14** |
| **PN96_RS12950** | | **GGDEF and EAL domain-containing protein** | **-4.07** |
| PN96_RS15200 | | polysaccharide biosynthesis/export family protein | -4.07 |
| **PN96_RS06415** | | **aspartate/glutamate racemase family protein** | **-4.04** |
| PN96_RS17225 | | ABC transporter substrate-binding protein | -4.01 |
| **PN96_RS23280** | | **CPBP family intramembrane glutamic endopeptidase** | **-4.00** |
| **PN96_RS23260** | ***alr*** | **alanine racemase** | **-3.94** |
| PN96_RS15550 | | L-lactate permease | -3.93 |
| PN96_RS15230 | | glycosyltransferase | -3.92 |
| PN96_RS15195 | | outer membrane β-barrel protein | -3.89 |
| PN96_RS21480 | | D-(-)-3-hydroxybutyrate oligomer hydrolase | -3.89 |
| **PN96_RS06420** | ***puuE*** | **allantoinase PuuE** | **-3.86** |
| PN96_RS03275 | | OmpA family protein | -3.79 |
| PN96_RS15210 | | putative capsular polysaccharide synthesis family protein | -3.79 |
| PN96_RS06390 | | alanine--glyoxylate aminotransferase family protein | -3.76 |
| PN96_RS15555 | *lldD* | FMN-dependent L-lactate dehydrogenase LldD | -3.72 |
| PN96_RS06695 | | EthD domain-containing protein | -3.67 |
| PN96_RS04705 | | O-antigen ligase family protein | -3.67 |
| PN96_RS04695 | | glycosyltransferase | -3.67 |
| PN96_RS03060 | | hypothetical protein | -3.66 |
| PN96_RS17420 | | hypothetical protein | -3.64 |
| **PN96_RS06385** | | **allantoate amidohydrolase** | **-3.62** |
| PN96_RS15220 | | O-antigen ligase family protein | -3.56 |
| PN96_RS19355 | *betA* | choline dehydrogenase | -3.54 |
| PN96_RS15205 | | polysaccharide biosynthesis tyrosine autokinase | -3.53 |
| PN96_RS21310 | | glycosyl transferase family protein | -3.52 |
| PN96_RS15225 | | putative capsular polysaccharide synthesis family protein | -3.52 |
| PN96_RS16085 | | hypothetical protein | -3.50 |
| PN96_RS15190 | | undecaprenyl-phosphate glucose phosphotransferase | -3.50 |
| PN96_RS00590 | *arcA* | arginine deiminase | -3.47 |
| **PN96_RS06380** | | **DUF5718 family protein** | **-3.46** |

| | | | |
|---|---|---|---|
| PN96_RS07915 | | hypothetical protein | -3.46 |
| PN96_RS17845 | | hypothetical protein | -3.46 |
| PN96_RS15215 | | glycosyltransferase | -3.45 |
| PN96_RS17415 | | LruC domain-containing protein | -3.45 |
| PN96_RS15235 | | oligosaccharide flippase family protein | -3.44 |
| PN96_RS06700 | | MFS transporter | -3.43 |
| PN96_RS06275 | *ureA* | urease subunit gamma | -3.41 |
| PN96_RS22025 | | TetR/AcrR family transcriptional regulator | -3.38 |
| PN96_RS17230 | | branched-chain amino acid ABC transporter permease | -3.38 |
| PN96_RS22585 | | hypothetical protein | -3.34 |
| PN96_RS17065 | | porin | -3.32 |
| **PN96_RS22685** | | **dipeptidase** | **-3.26** |
| PN96_RS16325 | | ABC transporter ATP-binding protein | -3.23 |
| PN96_RS06245 | | HupE/UreJ family protein | -3.23 |
| PN96_RS17525 | | DUF134 domain-containing protein | -3.22 |
| PN96_RS06255 | | urease accessory UreF family protein | -3.22 |
| PN96_RS17530 | | NifB/NifX family Fe-Mo cluster-binding protein | -3.21 |
| PN96_RS05970 | | DUF3305 domain-containing protein | -3.19 |
| PN96_RS16355 | *ahpC* | alkyl hydroperoxide reductase subunit C | -3.19 |
| PN96_RS05955 | | molecular chaperone | -3.15 |
| PN96_RS06265 | *ureC* | urease α-subunit | -3.14 |
| PN96_RS05105 | | EamA family transporter | -3.14 |
| PN96_RS06250 | *ureG* | urease accessory protein UreG | -3.13 |
| PN96_RS17535 | *nqrM* | (Na+)-NQR maturation NqrM | -3.13 |
| PN96_RS06285 | *urtE* | urea ABC transporter ATP-binding subunit UrtE | -3.13 |
| PN96_RS06260 | *ureE* | urease accessory protein UreE | -3.12 |
| PN96_RS05965 | | DUF3306 domain-containing protein | -3.12 |
| PN96_RS22995 | | glutaredoxin | -3.11 |
| PN96_RS09125 | | TRAP transporter substrate-binding protein | -3.09 |
| PN96_RS06280 | | urease accessory protein UreD | -3.07 |
| PN96_RS06270 | | urease β-subunit | -3.06 |
| PN96_RS17540 | | hypothetical protein | -3.06 |
| PN96_RS15240 | | VanZ family protein | -3.06 |
| PN96_RS09120 | | TRAP transporter small permease | -3.03 |
| PN96_RS13150 | | prepilin peptidase | -2.99 |
| PN96_RS06290 | *urtD* | urea ABC transporter ATP-binding protein UrtD | -2.98 |
| PN96_RS06540 | *hcaC* | 3-phenylpropionate/cinnamic acid dioxygenase ferredoxin subunit | -2.98 |
| PN96_RS09670 | *flgK* | flagellar hook-associated protein FlgK | -2.98 |
| PN96_RS12720 | | energy transducer TonB | -2.95 |
| PN96_RS12695 | | TonB-dependent receptor | -2.87 |
| PN96_RS16395 | | 1-pyrroline-5-carboxylate dehydrogenase | -2.87 |
| PN96_RS09750 | | FlgO family outer membrane protein | -2.86 |
| PN96_RS05960 | | 4Fe-4S dicluster domain-containing protein | -2.86 |
| PN96_RS05950 | | transcriptional initiation protein Tat | -2.85 |

| | | | |
|---|---|---|---|
| PN96_RS09675 | *flgJ* | flagellar assembly peptidoglycan hydrolase FlgJ | -2.84 |
| **PN96_RS12930** | ***glnA*** | **glutamate--ammonia ligase** | **-2.84** |
| PN96_RS02605 | | flagellar protein FliT | -2.83 |
| PN96_RS18640 | | Lrp/AsnC family transcriptional regulator | -2.82 |
| PN96_RS04690 | | chain-length determining protein | -2.82 |
| PN96_RS15470 | | hypothetical protein | -2.80 |
| PN96_RS13415 | *fadB* | fatty acid oxidation complex α-subunit FadB | -2.79 |
| PN96_RS12705 | | MotA/TolQ/ExbB proton channel family protein | -2.78 |
| PN96_RS09660 | | flagellin | -2.78 |
| PN96_RS12715 | | biopolymer transporter ExbD | -2.77 |
| **PN96_RS16390** | ***putP*** | **sodium/proline symporter PutP** | **-2.76** |
| PN96_RS12710 | | MotA/TolQ/ExbB proton channel family protein | -2.76 |
| PN96_RS06295 | *urtC* | urea ABC transporter permease subunit UrtC | -2.75 |
| PN96_RS16360 | *ahpF* | alkyl hydroperoxide reductase subunit F | -2.74 |
| PN96_RS12700 | | DUF3450 domain-containing protein | -2.74 |
| PN96_RS19795 | | transporter substrate-binding domain-containing protein | -2.71 |
| PN96_RS13420 | *fadA* | acetyl-CoA C-acyltransferase FadA | -2.71 |
| **PN96_RS19725** | ***mqo*** | **malate dehydrogenase (quinone)** | **-2.70** |
| **PN96_RS16400** | ***putA*** | **bifunctional proline dehydrogenase/L-glutamate gamma-semialdehyde dehydrogenase PutA** | **-2.69** |
| PN96_RS24345 | | ABC transporter ATP-binding protein | -2.67 |
| PN96_RS09745 | *flgP* | flagellar assembly lipoprotein FlgP | -2.66 |
| PN96_RS06300 | *urtB* | urea ABC transporter permease subunit UrtB | -2.64 |
| **PN96_RS06375** | | **chromate transporter** | **-2.63** |
| PN96_RS04750 | | SLBB domain-containing protein | -2.62 |
| PN96_RS08175 | | hypothetical protein | -2.62 |
| PN96_RS12725 | | hypothetical protein | -2.61 |
| PN96_RS17240 | | amidohydrolase family protein | -2.58 |
| PN96_RS20765 | | hypothetical protein | -2.58 |
| PN96_RS09680 | | flagellar basal body P-ring protein FlgI | -2.58 |
| PN96_RS19800 | | response regulator | -2.57 |
| PN96_RS06535 | *hcaF* | phenylpropionate/cinnamic acid dioxygenase β-subunit | -2.56 |
| PN96_RS19850 | | MFS transporter | -2.56 |
| PN96_RS19370 | *choV* | choline ABC transporter ATP-binding protein | -2.54 |
| PN96_RS18285 | | SDR family oxidoreductase | -2.53 |
| PN96_RS24340 | | branched-chain amino acid ABC transporter ATP-binding protein/permease | -2.52 |
| PN96_RS05940 | *fdh3B* | formate dehydrogenase FDH3 β-subunit | -2.52 |
| PN96_RS05945 | | formate dehydrogenase α-subunit | -2.52 |
| **PN96_RS03780** | | **circularly permuted type 2 ATP-grasp protein** | **-2.50** |
| PN96_RS07415 | | cysteine desulfurase-like protein | -2.50 |
| PN96_RS07420 | | 2OG-Fe dioxygenase family protein | -2.47 |
| PN96_RS20430 | | TRAP transporter small permease subunit | -2.44 |
| PN96_RS19790 | | ATP-binding protein | -2.44 |

| | | | |
|---|---|---|---|
| PN96_RS06305 | *urtA* | urea ABC transporter substrate-binding protein | -2.43 |
| PN96_RS22900 | | OmpA family protein | -2.42 |
| PN96_RS02600 | *fliD* | flagellar filament capping protein FliD | -2.42 |
| PN96_RS05935 | | formate dehydrogenase subunit gamma | -2.40 |
| PN96_RS19365 | *choW* | choline ABC transporter permease subunit | -2.39 |
| **PN96_RS13145** | ***cpaB*** | **Flp pilus assembly protein CpaB** | **-2.38** |
| PN96_RS17935 | | SDR family oxidoreductase | -2.35 |
| PN96_RS03995 | | calcium-binding protein | -2.35 |
| PN96_RS09685 | *flgH* | flagellar basal body L-ring protein FlgH | -2.34 |
| PN96_RS09690 | *flgG* | flagellar basal-body rod protein FlgG | -2.33 |
| **PN96_RS23255** | | **sodium:proton antiporter** | **-2.32** |
| PN96_RS15025 | | redoxin family protein | -2.26 |
| PN96_RS22035 | | APC family permease | -2.22 |
| PN96_RS20665 | | DUF1800 domain-containing protein | -2.21 |
| PN96_RS24035 | | IS3 family transposase - programmed frameshift | -2.21 |
| PN96_RS02610 | *fliS* | flagellar export chaperone FliS | -2.20 |
| PN96_RS18290 | | acetyl-CoA C-acetyltransferase | -2.20 |
| PN96_RS11990 | *cysD* | sulfate adenylyltransferase subunit CysD | -2.19 |
| PN96_RS16300 | *tauA* | taurine ABC transporter substrate-binding protein | -2.18 |
| PN96_RS15350 | | type I secretion system permease/ATPase | -2.17 |
| PN96_RS09695 | | flagellar basal body rod protein FlgF | -2.17 |
| PN96_RS22120 | *fdrA* | acyl-CoA synthetase FdrA | -2.17 |
| PN96_RS18345 | | hypothetical protein | -2.15 |
| PN96_RS17220 | | membrane protein | -2.13 |
| PN96_RS18645 | | ornithine cyclodeaminase | -2.12 |
| PN96_RS23040 | | methyl-accepting chemotaxis protein | -2.12 |
| PN96_RS00935 | | LysE/ArgO family amino acid transporter | -2.12 |
| PN96_RS09755 | | flagellar assembly protein FlgT | -2.11 |
| PN96_RS15355 | | calcium-binding protein | -2.11 |
| PN96_RS03915 | | septation protein A | -2.09 |
| PN96_RS22785 | | 4a-hydroxytetrahydrobiopterin dehydratase | -2.07 |
| PN96_RS09665 | *flgL* | flagellar hook-associated protein FlgL | -2.06 |
| PN96_RS16290 | | ABC transporter permease subunit | -2.06 |
| PN96_RS11985 | *cysN* | sulfate adenylyltransferase subunit CysN | -2.04 |
| PN96_RS05385 | | aldehyde dehydrogenase family protein | -2.04 |
| PN96_RS02595 | *flaG* | flagellar protein FlaG | -2.04 |
| PN96_RS19540 | | ExeM/NucH family extracellular endonuclease | -2.02 |
| PN96_RS15360 | | LuxR C-terminal-related transcriptional regulator | -2.00 |
| **Upregulated genes in Δ*ntrBC*** | | | |
| PN96_RS11665 | | lysophospholipid acyltransferase family protein | 7.73 |
| PN96_RS13625 | *punC* | purine nucleoside transporter PunC | 7.38 |
| PN96_RS09210 | | isochorismatase family protein | 7.15 |
| PN96_RS15300 | | aromatic ring-hydroxylating dioxygenase α-subunit | 7.03 |
| PN96_RS16160 | | SulP family inorganic anion transporter | 6.96 |

| | | | |
|---|---|---|---|
| PN96_RS07115 | | ATP-binding protein | 6.57 |
| PN96_RS08155 | | tRNA-Ser | 2.80 |
| PN96_RS08160 | | L-alanine exporter AlaE | 2.76 |
| PN96_RS17795 | | ribosome recycling factor family protein | 2.20 |
| PN96_RS14570 | *frdD* | fumarate reductase subunit FrdD | 2.15 |
| PN96_RS11245 | *gltB* | glutamate synthase large subunit | 2.01 |
| PN96_RS14580 | | succinate dehydrogenase/fumarate reductase Fe-S subunit | 2.00 |

[a]Bold, genes directly regulated by NtrC as determined by ChIP-seq

**Table S3. Differentially expressed genes in nitrogen-starved Δ*nifA* mutant versus the parent**

| Locus tag | Gene | Description | $Log_2$ fold-change |
|---|---|---|---|
| **Downregulated genes in Δ*nifA*** | | | |
| PN96_RS19445 | *nifD* | nitrogenase Fe-Mo protein α-subunit | -10.69 |
| PN96_RS19440 | *nifH* | nitrogenase Fe protein | -10.51 |
| PN96_RS19450 | *nifK* | nitrogenase Fe-Mo protein β-subunit | -10.34 |
| PN96_RS19455 | *nifT* | putative nitrogen fixation protein NifT | -10.22 |
| PN96_RS19490 | *nifU* | Fe-S cluster assembly protein NifU | -10.16 |
| PN96_RS19620 | *rnfA* | RnfABCDGE type electron transport complex subunit A | -10.06 |
| PN96_RS19495 | *nifS* | cysteine desulfurase NifS | -10.05 |
| PN96_RS19625 | *rnfB* | RnfABCDGE type electron transport complex subunit B | -9.76 |
| PN96_RS19500 | *nifV* | homocitrate synthase | -9.69 |
| PN96_RS19460 | | NifB/NifX family Fe-Mo cluster-binding protein | -9.69 |
| PN96_RS19465 | | hypothetical protein | -9.62 |
| PN96_RS19630 | *rnfC* | RnfABCDGE type electron transport complex subunit C | -9.56 |
| PN96_RS19505 | *nifW* | nitrogenase-stabilizing/protective protein NifW | -9.48 |
| PN96_RS19510 | *nifZ* | nitrogen fixation protein NifZ | -9.30 |
| PN96_RS19515 | *nifM* | peptidylprolyl isomerase | -9.19 |
| PN96_RS19520 | *modA1* | molybdate ABC transporter substrate-binding protein | -9.14 |
| PN96_RS24200 | | hypothetical protein | -9.06 |
| PN96_RS19635 | *rnfD* | RnfABCDGE type electron transport complex subunit D | -9.01 |
| PN96_RS19470 | *nifE* | nitrogenase Fe-Mo cofactor biosynthesis protein NifE | -8.98 |
| PN96_RS19525 | *modB1* | molybdate ABC transporter permease subunit | -8.78 |
| PN96_RS19640 | *rnfG* | RnfABCDGE type electron transport complex subunit G | -8.70 |
| PN96_RS19530 | *modC1* | molybdenum ABC transporter ATP-binding protein | -8.68 |
| PN96_RS19480 | | NifB/NifX family Fe-Mo cluster-binding protein | -8.62 |
| PN96_RS19475 | *nifN* | nitrogenase Fe-Mo cofactor biosynthesis protein NifN | -8.58 |
| PN96_RS19485 | *fdxB* | ferredoxin III, nif-specific | -8.51 |
| PN96_RS19645 | *rnfE* | RnfABCDGE type electron transport complex subunit E | -8.50 |
| PN96_RS19650 | *rnfH* | RnfH family protein | -8.09 |
| PN96_RS19655 | *nifF* | flavodoxin | -7.99 |
| PN96_RS19610 | *nifA* | nif-specific transcriptional activator NifA | -5.13 |
| PN96_RS07685 | | hybrid-cluster NAD(P)-dependent oxidoreductase | -4.82 |
| PN96_RS07680 | *hcp* | hydroxylamine reductase | -4.30 |
| PN96_RS19605 | *nifB* | nitrogenase cofactor biosynthesis protein NifB | -2.83 |
| PN96_RS16355 | *ahpC* | alkyl hydroperoxide reductase subunit C | -2.79 |
| PN96_RS19595 | *nifQ* | nitrogen fixation protein NifQ | -2.76 |
| PN96_RS17525 | | DUF134 domain-containing protein | -2.74 |
| PN96_RS17530 | | NifB/NifX family Fe-Mo cluster-binding protein | -2.72 |
| PN96_RS19550 | | DMT family transporter | -2.71 |
| PN96_RS15025 | | redoxin family protein | -2.66 |
| PN96_RS17535 | *nqrM* | (Na+)-NQR maturation NqrM | -2.58 |
| PN96_RS19660 | | hypothetical protein | -2.45 |

| | | | |
|---|---|---|---|
| PN96_RS17540 | | hypothetical protein | -2.44 |
| PN96_RS16360 | *ahpF* | alkyl hydroperoxide reductase subunit F | -2.40 |
| PN96_RS24035 | | IS3 family transposase - programmed frameshift | -2.26 |
| PN96_RS04695 | | glycosyltransferase | -2.23 |
| PN96_RS19540 | | ExeM/NucH family extracellular endonuclease | -2.00 |

### Table S4. Bacterial strains

| Strain/ designation | Derived from | Relevant genotype[a] | Comments | Source |
|---|---|---|---|---|
| TND1964/ wild-type | ATCC 14048 | pMMB-*tfoX* (Vc) | | [1] |
| NWH003/ Parent | TND1964 | Δ*dns*::$Km^R$ | | This study |
| NWH004/ Parent | TND1964 | Δ*dns*::$Sp^R$ | | This study |
| NWH005 | TND1964 | Δ*ntrBC* Δ*dns*::$Sp^R$ | Intermediate used to make NWH014, 043, and 064 | This study |
| NWH006 | TND1964 | Δ*dns*::$Km^R$-*lacZ* | Intermediate used to make NWH010 and 017 | This study |
| NWH010/ Parent | NWH006 | Δ*dns*::$Sp^R$-$P_{nifLA}$-*lacZ* | Parent with $P_{nifLA}$-LacZ reporter | This study |
| NWH014/ Δ*ntrBC* | NWH005 | Δ*ntrBC,* Δ*dns*::$Km^R$ | | This study |
| NWH015/ Δ*ntrBC* | NWH014 | Δ*ntrBC* Δ*dns*::$Sp^R$-$P_{nifLA}$-*lacZ* | Δ*ntrBC* mutant with $P_{nifLA}$-LacZ reporter | This study |
| NWH017 | NWH006 | Δ*dns*::$Sp^R$-*lacZ* | Promoterless LacZ control | This study |
| NWH036/ Δ*nifA* | TND1964 | Δ*nifA,* Δ*dns*::$Km^R$ | | This study |
| NWH037/ Δ*nifL* | TND1964 | Δ*nifL,* Δ*dns*::$Km^R$ | | This study |
| NWH038/ Δ*rnF* | TND1964 | Δ*rnf,* Δ*dns*::$Km^R$ | *rnf* operon deletion (*RS19650-19620*) | This study |
| NWH039/ Δ*nifF* | TND1964 | Δ*nifF,* Δ*dns*::$Km^R$ | | This study |
| NWH042/ Δ*glnB* | TND1964 | Δ*glnB,* Δ*dns*::$Km^R$ | | This study |
| NWH043/ Δ*ntrBC*Δ*nifA* | NWH005 | Δ*ntrBC,* Δ*nifA,* Δ*dns*::$Km^R$ | | This study |
| NWH044/ Parent | NWH006 | Δ*dns*::$Sp^R$-$P_{nifHDK}$-*lacZ* | Parent with $P_{nifHDK}$-LacZ reporter | This study |
| NWH047/ Δ*glnK* | TND1964 | Δ*glnK,* Δ*dns*::$Km^R$ | | This study |
| NWH050/ ΔPII | NWH042 | Δ*glnB,* Δ*glnK,* Δ*dns*::$Sp^R$ | | This study |
| NWH060 | TND1964 | Δ*dns*::$Km^R$-$P_{ntrBC}$-*ntrBC* | Intermediate used to make NWH064 | This study |
| NWH061 | NWH064 | Δ*ntrBC,* Δ*dns*::$Sp^R$-$P_{glnA}$-*ntrBC* | Δ*ntrBC* complemented. Untagged ChIP-seq control | This study |
| NWH064 | NWH005 | Δ*ntrBC,* Δ*dns*::$Km^R$-$P_{ntrBC}$-*ntrBC* | Intermediate used to make NWH061 | This study |
| NWH066 | TND1964 | Δ*dns*::$Sp^R$-$P_{glnA}$-*ntrB*-*ntrC*::His6x | Intermediate used to make NWH071 | This study |
| NWH071 | NWH014 | Δ*ntrBC,* Δ*dns*::$Sp^R$-$P_{glnA}$-*ntrB*-*ntrC*::His6x | Δ*ntrBC* complemented with WT *ntrB* and a C-terminal His6x-tagged *ntrC* for ChIP-seq | This study |
| NWH090/ Δ*nifA* | NWH036 | Δ*nifA* Δ*dns*::$Sp^R$-$P_{nifLA}$-*lacZ* | Δ*nifA* mutant with $P_{nifLA}$-LacZ reporter | This study |
| NWH105/ parent | TND1964 | Δ*dns*::$Sp^R$-$P_{tet}$-*nifA*-$P_{lacI}$-*tetR* | Parent with aTc-inducible *nifA* | This study |
| NWH106/ Δ*ntrBC* | NWH014 | Δ*ntrBC,* Δ*dns*::$Sp^R$-$P_{tet}$-*nifA*-$P_{lacI}$-*tetR* | Δ*ntrBC* mutant with aTc-inducible *nifA* | This study |
| NWH108/ Δ*ntrBC*Δ*nifA* | NWH043 | Δ*ntrBC,* Δ*nifA* Δ*dns*::$Sp^R$-$P_{tet}$-*nifA*-$P_{lacI}$-*tetR* | Δ*ntrBC*Δ*nifA* mutant with aTc-inducible *nifA* | This study |
| NWH111 ΔPIIΔ*ntrBC* | NWH042 | Δ*glnB,* Δ*glnK,* Δ*ntrBC,* Δ*dns*::$Sp^R$ | | This study |
| NWH112 ΔPIIΔ*nifA* | NWH042 | Δ*glnB,* Δ*glnK,* Δ*nifA,* Δ*dns*::$Sp^R$ | | This study |
| NWH126/ Δ*ntrBC*Δ*nifA* | NWH043 | Δ*ntrBC,* Δ*nifA,* Δ*dns*::$Sp^R$-$P_{nifLA}$-*lacZ* | Δ*ntrBC*Δ*nifA* mutant with $P_{nifLA}$-LacZ reporter | This study |

| | | | | |
|---|---|---|---|---|
| NWH127/ Δ*ntrBC*Δ*nifA,* $P_{tet}$-*nifA* | NWH043 | Δ*ntrBC,* Δ*nifA,* Δ*dns*::$Sp^R$-$P_{nifLA}$-*lacZ*-$P_{tet}$-*nifA*-$P_{lacI}$-*tetR* | Δ*ntrBC*Δ*nifA* mutant with $P_{nifLA}$-LacZ reporter and aTc-inducible *nifA* | This study |

[a]All strains retain the pMMB-*tfoX* (Vc) plasmid following cloning, but the plasmid was omitted from other rows for brevity. All strains are thus resistance to carbenicillin due to the resistance cassette on the plasmid.

### Table S5. Primers

| Primer | Sequence (5'→ 3')[a] | Description[b] |
|---|---|---|
| NH014 | CTCTGCACCACTACCGTC | F; Δ*dns*::XXX screening |
| NH015 | CGAATACCGATGTCGCTGC | R; Δ*dns*::XXX screening |
| NH022 | CGAGGTGAAGATCATTCATTTCC | F; universal primer; upstream for Δ*dns*::XXX |
| NH025 | CTTAGTGATTGGGTCACTCATTGG | R; universal primer; downstream for Δ*dns*::XXX |
| NH028 | CTATGGACACGGGTAAAATCATACC | R; Δ*dns*::Sp$^R$ internal for screening |
| NH031 | CTCGATGTCGTTCGAGTCC | F; upstream for Δ*ntrBC*; SOE |
| NH032 | aatcaaagtgtcagtacaactccttCACATTATGTCCTTGTTTGCG | R; upstream for Δ*ntrBC* |
| NH033 | ttaacgcaaacaaggacataatgtgAAGGAGTTGTACTGACACTTTG | F; downstream for Δ*ntrBC* |
| NH034 | GATCACCCAGTACCGCTAC | R; downstream for Δ*ntrBC*; SOE |
| NH035 | GAGGGACTCACACTCTTTCAG | F; Δ*ntrBC* screening |
| NH036 | CCGACAGTTGCTTATCACTG | R; Δ*ntrBC* screening |
| NH041 | ctgcaggtcgacggatccccggaatCACACAGGAAACAGCTATGACCATG | F; pHRP309 *lacZ* + RBS in Δ*dns*::Km$^R$-*lacZ* |
| NH042 | tttaactatgaaatacctgttctctCTGTCGCACCTGCTTGATCG | R; pHRP309 *lacZ* + RBS in Δ*dns*::Km$^R$-*lacZ* |
| NH043 | cgattgtcgcgattggtgaggattaTGTAGGCTGGAGCTGCTTC | F; For Km$^R$ in Δ*dns*::Km$^R$-*lacZ* |
| NH044 | catggtcatagctgtttcctgtgtgATTCCGGGGATCCGTCGAC | R; For Km$^R$ in Δ*dns*::Km$^R$-*lacZ* |
| NH045 | aacttcgaagcagctccagcctacaTAATCCTCACCAATCGCGAC | R; upstream of *dns* for Δ*dns*::Km$^R$-*lacZ* |
| NH046 | ggccgcgatcaagcaggtgcgacagAGAGAACAGGTATTTCATAGTTAAAGTC | F; downstream for of *dns* for Δ*dns*::Km$^R$-*lacZ* |
| NH049 | CGCTTGCGTTTTCCCTTG | F; Δ*dns*::Sp$^R$ internal for screening |
| NH053 | CAGACCATTTTCAATCCGCAC | R; universal SOE for P$_{xxx}$-*lacZ* fusions; 5' in *lacZ* |
| NH054 | tggcatacgctatgcaccaccttcgCACTTTATGCTTCCGACCTG | R; upstream for Δ*dns*::Sp$^R$-P$_{nifLA}$-*lacZ*, Sp$^R$-*P*$_{nifLA}$ junction |
| NH055 | ggctgcaggtcggaagcataaagtgCGAAGGTGGTGCATAGCG | F; for Δ*dns*::Sp$^R$-P$_{nifLA}$-*lacZ*, Sp$^R$-*P*$_{nifLA}$ junction; Δ*nifL* screening |
| NH056 | catggtcatagctgtttcctgtgtgCGTTTGGCGATATAAGACCGTC | R; for Δ*dns*::Sp$^R$-P$_{nifLA}$-*lacZ*, *P*$_{nifLA}$-*lacZ* junction |
| NH057 | atcgacggtcttatatcgccaaacgCACACAGGAAACAGCTATGACCATG | F; for Δ*dns*::Sp$^R$-P$_{nifLA}$-*lacZ*, *lacZ* downstream arm |
| NH058 | GGCTGAATATCGACGGTTTCC | F; Δ*dns*::*lacZ* internal screening |
| NH064 | catggtcatagctgtttcctgtgtgCACTTTATGCTTCCGACCTG | R; to replace Km$^R$ with Sp$^R$ in Δ*dns*::Km$^R$-*lacZ* |
| NH065 | ggctgcaggtcggaagcataaagtgCACACAGGAAACAGCTATGACCATG | F; to replace Km$^R$ with Sp$^R$ in Δ*dns*::Km$^R$-*lacZ* |
| NH105 | CAGCTCTGGAATGTCTTG | F; upstream for Δ*rnf*; SOE |
| NH106 | gtcgacggatccccggaatCAATCCTTCGGACATAACAG | R; upstream for Δ*rnf* |
| NH107 | agaagcagctccagcctacaGATTAATTGTCACCTTTGCG | F; downstream for Δ*rnf* |
| NH108 | CGAGAACTAATCCAAAACTTC | R; downstream Δ*rnf*; SOE |
| NH109 | CGATATAAGACCGTCGATG | F; Δ*rnf* screening |
| NH110 | CGGTTTATCCGCAACCTC | R; Δ*rnf* screening |
| NH111 | GGTGCATGTGGATATGTTTG | F; upstream for Δ*nifF*; SOE |
| NH112 | gtcgacggatccccggaatGCTCATAATTAGCTCCTTGC | R; upstream for Δ*nifF* |
| NH113 | agaagcagctccagcctacaGAGCTATGAATAAAAATAAAGCTATTAAC | F; downstream for Δ*nifF* |
| NH114 | GACATTCATTATTACGAGTCGC | R; downstream for Δ*nifF*; SOE |
| NH115 | GATTAATTGTCACCTTTGCG | F; Δ*nifF* screening |

| | | |
|---|---|---|
| NH116 | CCTATTGTTCACCATTCC | R; Δ*nifF* screening |
| NH117 | agaagcagctccagcctacaAACGATCTCAGACAGCCGCC | F; downstream for Δ*nifL* |
| NH118 | CCGCCAATAACTGACGCTC | R; Δ*nifL* screening; $P_{nifLA}$-*lacZ*::$P_{tet}$-*nifA* screening |
| NH119 | CCAAGTGATACATTAATAGACG | F; upstream for Δ*glnB*; SOE |
| NH120 | gtcgacggatccccggaatCTTCATGTTCATCCCTTAAG | R; upstream for Δ*glnB* |
| NH121 | agaagcagctccagcctacaGCGATTTAACATTTCACTC | F; downstream for Δ*glnB* |
| NH122 | CTCCTCAGGAACAAGAAG | R; downstream for Δ*glnB*; SOE |
| NH123 | CGTCACTGCTGTCTGATG | F; Δ*glnB* screening |
| NH124 | CTCGATGCATTGAATATCTTCAG | R; Δ*glnB* screening |
| NH125 | gcatgcagagaagaaccctcCACTTTATGCTTCCGACCTG | R; upstream for Δ*dns*::$Sp^R$-$P_{nifHDK}$-*lacZ*, $Sp^R$-$P_{nifHDK}$ junction |
| NH126 | caggtcggaagcataaagtgGAGGGTTCTTCTCTGCATG | F; for Δ*dns*::$Sp^R$-$P_{nifHDK}$-*lacZ*, $Sp^R$-$P_{nifHDK}$ junction |
| NH127 | tcatagctgtttcctgtgtgTAATTTAGACTCATGACCGTTG | R; for Δ*dns*::$Sp^R$-$P_{nifHDK}$-*lacZ*, $P_{nifHDK}$-*lacZ* junction |
| NH128 | acggtcatgagtctaaattaCACACAGGAAACAGCTATGACCATG | F; for Δ*dns*::$Sp^R$-$P_{nifHDK}$-*lacZ*, *lacZ* downstream arm |
| NH129 | GATGAGCTTAAGACGCTC | F; upstream for Δ*glnK*; SOE |
| NH130 | gtcgacggatccccggaatCATTAAATTCTCTCCGAAGC | R; upstream for Δ*glnK* |
| NH131 | agaagcagctccagcctacaGCACTTTAAAAGTATTCAAGGG | F; downstream for Δ*glnK* |
| NH132 | CGAATATCCGGGAACCAG | R; downstream for *glnK*; SOE |
| NH133 | CTGTGAAAGCACCGCTAG | F; Δ*glnK* screening |
| NH134 | CGCCGTTATCAACGTACAT | R; Δ*glnK* screening |
| NH136 | ctgcaggtcgacggatccccggaatGACCGTAAACGGGGCAAAG | F; for Δ*dns*::$Km^R$-$P_{ntrBC}$-*ntrBC*, $Km^R$-$P_{ntrBC}$ |
| NH137 | tttaactatgaaatacctgttctctGTCTAATCACAAAAGGGTGGTG | R; for Δ*dns*::$Km^R$-$P_{ntrBC}$-*ntrBC*, *ntrBC*::Δ*dns* junction |
| NH138 | gcgcaccaccccttttgtgattagacAGAGAACAGGTATTTCATAGTTAAAGTC | F; for Δ*dns*::$Km^R$-$P_{ntrBC}$-*ntrBC*, *ntrBC*::Δ*dns* downstream arm |
| NH139 | ctgcaggtcgacggatccccggaatCTCTCTGGCGACATTTGC | F; for replacing $Km^R$-$P_{ntrBC}$ with $Sp^R$-$P_{glnA}$ in Δ*dns*::$Km^R$-$P_{ntrBC}$-*ntrBC*; $Sp^R$-$P_{glnA}$ junction |
| NH140 | tccacattatgtccttgtttgcgttAAATAAGCGGTAAAAGCTCG | R; for replacing $Km^R$-$P_{ntrBC}$ with $Sp^R$-$P_{glnA}$ in Δ*dns*::$Km^R$-$P_{ntrBC}$-*ntrBC*; $P_{glnA}$-*ntrB* junction |
| NH141 | taaatcgagcttttaccgcttatttAACGCAAACAAGGACATAATG | F; for replacing $Km^R$-$P_{ntrBC}$ with $Sp^R$-$P_{glnA}$ in Δ*dns*::$Km^R$-$P_{ntrBC}$-*ntrBC*; $P_{glnA}$-*ntrB* junction |
| NH142 | GCTAAACTTGGTCAGATCG | R; for replacing $Km^R$-$P_{ntrBC}$ with $Sp^R$-$P_{glnA}$ in Δ*dns*::$Km^R$-$P_{ntrBC}$-*ntrBC*; downstream arm |
| NH147 | tggtggtgtgaacctgaaccGTACAACTCCTTTAATTTTCTGGTG | R; for Δ*dns*::$Sp^R$-$P_{glnA}$-*ntrB*-*ntrC*::His6x; *ntrC*-Hisx6 junction |
| NH148 | GGTTCAGGTTCACACCACCACCACCACCAC | F; linker + Hisx6 tag for C-terminal NtrC fusion |
| NH149 | GTGGTGGTGGTGGTGGTGTGAACCTGAACC | R; linker + Hisx6 tag for C-terminal NtrC fusion |
| NH150 | cacaccaccaccaccaccacTGACACTTTGATTAATAATACAGCATAGC | F; for Δ*dns*::$Sp^R$-$P_{glnA}$-*ntrB*-*ntrC*::His6x; linker +Hisx6 downstream arm |
| NH222 | CTAGTATTTCCCCTCTTTCTCTAG | R; upstream for $P_{tet}$-*nifA*-$P_{lacI}$-*tetR*; $P_{tet}$-*nifA* junction |

| | | |
|---|---|---|
| NH223 | CTCGGTACCAAATTCCAGAAAAG | F; downstream for $P_{tet}$-*nifA*-$P_{lacI}$-*tetR*; -*nifA*-$P_{lacI}$ junction |
| NH224 | actagagaaagaggggaaatactagATGATAGAAGATCATTCTCTTTTAGATTTAGAG | F; for $P_{tet}$-*nifA*-$P_{lacI}$-*tetR*; $P_{tet}$-*nifA* junction |
| NH225 | ctcttttctggaatttggtaccgagTCAGATTTGTTTCATCTCGATATT | R; for $P_{tet}$-*nifA*-$P_{lacI}$-*tetR*; *nifA*-$P_{lacI}$ junction |
| NH235 | tggttgtgtctcaaaatctctgatgCCTGTTCTCTCTGTCGCAC | R; upstream for Δ*dns*::$Sp^R$-$P_{nifLA}$-*lacZ*-$P_{tet}$-*nifA*-$P_{lacI}$-*tetR*; *nifA*-$P_{lacI}$; *lacZ*-$P_{tet}$ junction |
| NH236 | CATCAGAGATTTTGAGACACAACC | F; downstream for Δ*dns*::$Sp^R$-$P_{nifLA}$-*lacZ*- $P_{tet}$-*nifA*-$P_{lacI}$-*tetR*; *nifA*-$P_{lacI}$; *lacZ*-$P_{tet}$ junction |
| NH237 | CGGTCGCTACCATTACCAG | F; $P_{nifLA}$-*lacZ*::$P_{tet}$-*nifA* internal screening |
| SB001 | GGACATGATTCACTCCTAAACCG | F; upstream for Δ*nifA*; SOE |
| JL005 | gtcgacggatccccggaatCCGCCAATAACTGACGCTC | R; upstream for Δ*nifA* |
| JL006 | gaagcagctccagcctacaCAAGGCTGCTCAATATGACGC | F; downstream for Δ*nifA* |
| JL007 | GAGATAAGTGACGCAAGCCG | R; downstream for Δ*nifA*; SOE |
| JL008 | AACTGTTGCTCCTTCTCAGCG | F; upstream for Δ*nifL*; SOE |
| JL011 | TGGTTTGTCTGCTCTGTCGC | R; downstream for Δ*nifL*; SOE |
| JL016 | gtcgacggatccccggaatGTAATGCTGATGGCGAC | R; upstream for Δ*nifL* |
| JL049 | TCAGATTTGTTTCATCTCGATATT | R; Δ*nifA* screening |
| JL050 | ATGATAGAAGATCATTCTCTTTTAGATTTAGAG | F; Δ*nifA* screening |
| ABD123 | ATTCCGGGGATCCGTCGAC | F; universal FRT for $Sp^R$ or $Km^R$ |
| ABD124 | TGTAGGCTGGAGCTGCTTC | R; universal FRT for $Sp^R$ or $Km^R$ |
| miniFRT-F | ATTCCGGGGATCCGTCGACCTGCAGTTCAGAAGCAGCTCCAGCCTACA | F; Universal linker for deletion constructs |
| miniFRT-R | TGTAGGCTGGAGCTGCTTCTGAACTGCAGGTCGACGGATCCCCGGAAT | R; Universal linker for deletion constructs |

[a]Lowercase indicates overlapping regions for SOE PCR.

[b]F, forward; R, reverse
